# Solution architecture of G3BP1 reveals pH-dependent conformational switching underlying liquid-liquid phase separation

**DOI:** 10.1101/2025.03.27.645651

**Authors:** Xiao Han, Renhua Sun, Melissa A. Graewert, Qianyu Zhou, Tom Resink, Clement E. Blanchet, Gerald McInerney, Evren Alici, Hans-Gustaf Ljunggren, Marianne Farnebo, Dmitri Svergun, Adnane Achour

**Affiliations:** Science for Life Laboratory, Department of Medicine Solna, Karolinska Institute Solna, 17165, Sweden, and Division of Infectious Diseases, Karolinska University Hospital, Stockholm, 17177, Sweden; European Molecular Biology Laboratory EMBL, Hamburg Unit, c/o DESY, 22607, Hamburg, Germany; Department of Oncology and Pathology, Karolinska Institute, Stockholm, 17177, Sweden; Department of Microbiology, Tumor and Cell Biology, Karolinska Institute, Stockholm, 17165, Sweden; Centre for Hematology and Regenerative Medicine, Department of Medicine Huddinge, Karolinska Institute, Huddinge 14183, Sweden; Hematology Center, Karolinska University Hospital, Huddinge 14183, Sweden; Center for Infectious Medicine, Department of Medicine Huddinge, Karolinska Institute, Huddinge 14152, Sweden; BIOSAXS GmbH, c/o DESY, 22607, Hamburg, Germany

**Keywords:** Stress granules, G3BP1, pH, conformational dynamics, condensate, phase separation, intrinsically disordered region

## Abstract

G3BP1 is the central node and molecular switch in stress granule (SG) assembly. However, structural insights into full-length G3BP1 remain elusive owing to its extensive intrinsically disordered regions (IDRs). Using size-exclusion chromatography-coupled small-angle X-ray scattering (SEC-SAXS), we have characterized the solution architecture and conformational dynamics of full-length G3BP1. Under physiological conditions, G3BP1 adopts an elongated, head-to-head antiparallel homodimeric conformation, whereas acidification induces a pronounced conformational compaction. Subsequent biophysical studies reveal that this compact state enables robust RNA-mediated and, notably, homotypic phase separation *in vitro*. Deletion of the RGG region abolishes this acidity-induced compaction and markedly impairs phase separation, establishing a causal link between the RGG-dependent conformational switch and phase separation propensity. By moving beyond hypothetical models to experimental solution-state data, our work fills a longstanding void in the field and provides critical insights into the structural plasticity that underlies G3BP1 function, offering a missing structural link essential for deciphering the molecular mechanism of SG formation. We propose that stress-associated physicochemical changes, specifically localized acidification coupled with mRNA accumulation, trigger this reversible structural reconfiguration of G3BP1, thereby facilitating phase separation.

## Introduction

Eukaryotic cells harbor a wide range of dynamic membraneless organelles collectively referred to as ribonucleoprotein (RNP) granules^1–4^. Among these, stress granules (SGs) are cytoplasmic RNP assemblies formed via liquid-liquid phase separation (LLPS) in response to diverse physiological and pathological stresses, including hypoxia, viral infection, oxidative stress, osmotic stress, and heat-shock^5,6^. SGs play a crucial role in translational repression during stress and contribute to the regulation of mRNA stability, thereby preserving cellular homeostasis and promoting cell survival. These granules form within minutes of stress onset and disassemble rapidly once stress is relieved^7,8^, pointing to highly dynamic yet stringently controlled molecular mechanisms that mediate these transitions. The dysregulation of SGs has been implicated in a wide range of pathological contexts, including cancer, viral infections, and neurodegenerative diseases^9–14^. Notably, the aberrant persistence of SGs is increasingly recognized as a driver of chemoresistance and immunotherapy evasion across various malignancies, marking them as critical determinants of therapeutic failure^15–18^. Furthermore, the defective clearance of SGs is frequently observed in aging and neurodegenerative disorders, suggesting that impaired granule disassembly actively contributes to neurodegeneration^14,19,20^. Despite these well-established functional links, a comprehensive structural understanding of the key protein components remains elusive.

The GTPase-activating protein SH3 domain-binding proteins (G3BP1, G3BP-2a and G3BP-2b; collectively referred to as G3BPs) are core regulators of SG formation^21–25^. SG formation is initiated by the sudden release of mRNAs from polysomes upon translational arrest^26–29^. G3BP recognizes long, unfolded single-stranded RNA^22,29^, and acts as an RNA condenser, facilitating the intermolecular RNA-RNA interactions that stabilize RNP granules^30,31^. G3BP features an N-terminal nuclear transport factor 2-like (NTF2L) domain, followed by a highly acidic and negatively charged region (IDR1), a positively charged proline-rich domain (PxxP, IDR2), an RNA recognition motif (RRM), and a C-terminal arginine-glycine repeat (RGG) region (IDR3) (Fig. 1a). While structural studies have primarily focused on the NTF2L domain in complex with short linear motifs from viral and host partners^32–36^, the three distinct IDRs of G3BP1 are increasingly recognized as key regulators of phase separation^21–24^.

**Fig. 1:**
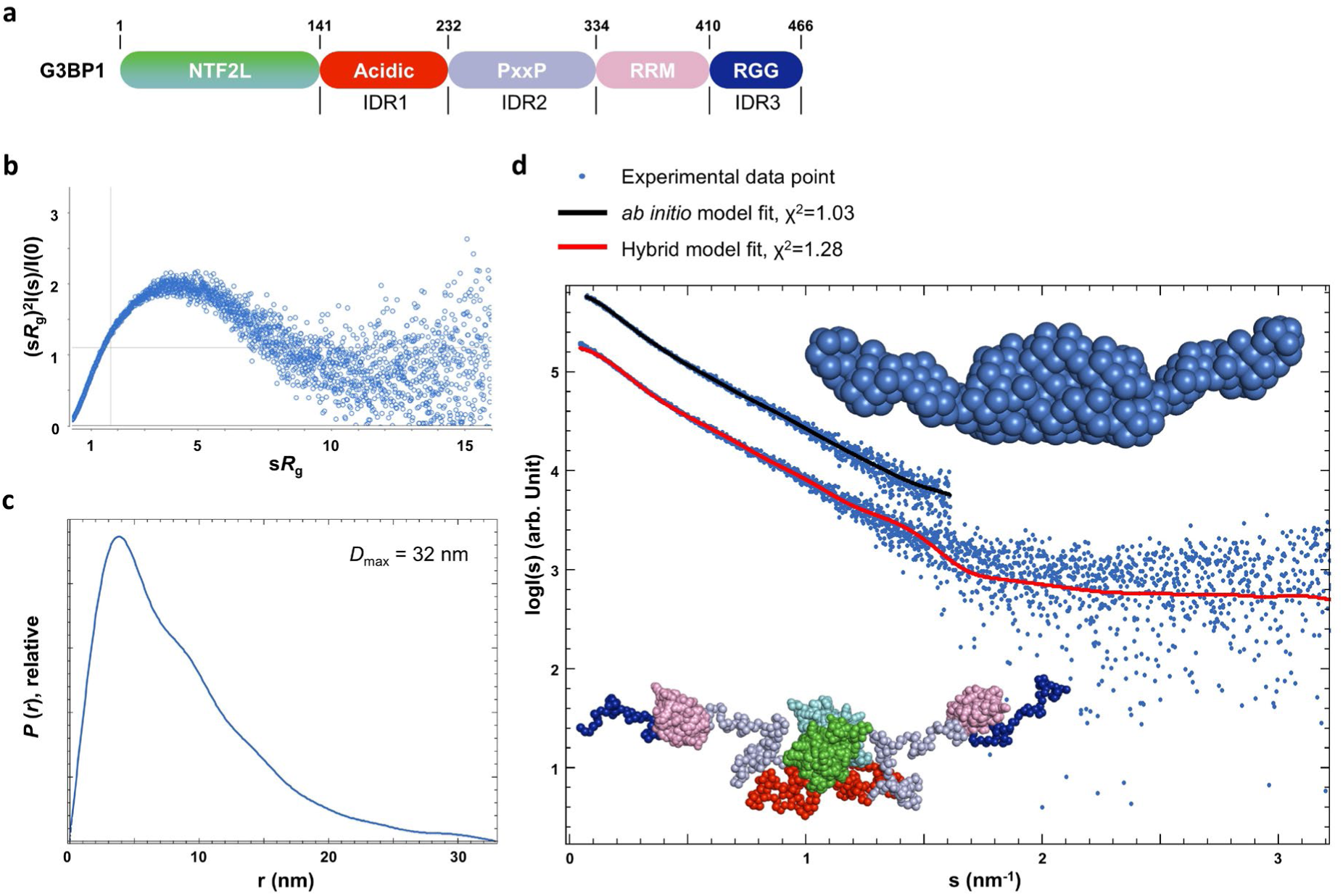
Full-length G3BP1 exhibits an elongated shape featuring constrained IDRs. **a**, Schematic diagram illustrating the domain organization of G3BP1. **b**, Experimental Scattering data of G3BP1 is shown as a normalized Kratky plot. **c**, Pair distance distribution function, *P(r)* of G3BP1. **d**, The *ab initio* model of G3BP1 (generated with DAMMIN) is presented in the upper inset as blue spheres. The black line describes the fit of the *ab initio* model to the experimental scattering data (plotted as log *I*(*s*) vs *s*, blue dots) within the restricted angular range. The hybrid model of G3BP1 (generated with CORAL) is presented in the lower inset as spheres, with the domains colored as indicated in (**a**). The red curve presents the fit of the hybrid model with experimental data. Curves have been shifted along the y-axis for better visibility. The buffer conditions for these analyses were 20 mM Tris-HCl (pH 7.5), 150 mM NaCl. The scattering data is displayed as a function of the momentum transfer s = 4π*sinθ/λ, where λ = 0.09464 nm is the wavelength and 2θ is the scattering angle.

Despite these advances, the overall architecture of full-length G3BP1 remains poorly defined. Due to its high IDR content, the protein is expected to exhibit substantial conformational heterogeneity. For instance, current models propose that G3BP1 transitions from a compact, autoinhibited state to an expanded, multivalent "open" conformation, initiating LLPS^22,23^. However, existing studies present an inconsistency: although both acidic pH and increased salt concentrations are shown to induce G3BP1 expansion, they lead to opposed functional outcomes - pH reduction facilitates condensation, whereas higher salinity promotes dissolution^22,23^. This discrepancy underscores the need for a systematic characterization of full-length G3BP1 to reconcile these observations and elucidate the molecular principles linking its conformational dynamics to phase separation capacity.

Here, we employed size-exclusion chromatography-coupled small-angle X-ray scattering (SEC-SAXS) and complementary biophysical assays, to define the solution architecture and conformational landscape of full-length G3BP1. Our results reveal that, under physiological conditions, G3BP1 adopts a more defined structural organization compared to AlphaFold2^37,38^ predictions and undergoes a pronounced conformational compaction upon acidification. Furthermore, we identify the RGG region as essential for this acidity-driven conformational switch and the phase separation propensity of G3BP1. These findings reconcile the structural plasticity of G3BP1 with its phase separation behavior, providing a structural framework for how environmental cues regulate stress granule assembly and establishing a molecular foundation for future therapeutic interventions targeting SG-associated pathologies.

## Results

### G3BP1 forms an elongated, head-to-head antiparallel homodimer featuring constrained IDRs

The primary structure and domain organization of G3BP1 reveal three IDRs, spanning the large acidic, PxxP, and RGG regions (Fig. 1a and Appendix Fig. S1a). While AlphaFold 2 predicts the structures of the NTF2L dimer and RRM domains with high confidence, the spatial relationship between these domains within the full-length G3BP1 remains ambiguous. The connecting linkers are modeled as low-confidence, flexible loops (Appendix Fig. S1b). Despite these IDRs, we successfully purified the recombinant full-length G3BP1. Dynamic light scattering (DLS) analysis revealed a dimer peak with a hydrodynamic radius (*R_h_*) of 8.2 ± 0.1 nm and a polydispersity index (PI) value of 0.2 ± 0.01, indicating a high degree of monodispersity (Appendix Fig. S1c).

We investigated the solution structure of G3BP1 using SEC-SAXS, a powerful technique to provide insights into the three-dimensional conformation and overall shape of biological macromolecules in solution. SEC-SAXS data was collected under physiological conditions (pH 7.5, 150 mM NaCl) (Fig. 1b). Analysis of the scattering profile yielded an estimated molecular weight of 119 kDa (Table 1), approximately doubling the theoretical monomeric weight of 52.5 kDa. This oligomeric state aligns with previous findings from multi-angle light scattering (MALS) and analytical ultracentrifugation (AUC), validating the constitutive dimerization of G3BP1^23^. Solution SEC-SAXS measurements revealed that G3BP1 adopts an elongated conformation, characterized by a maximum dimension (*D*_max_) of ∼32.4 nm and a radius of gyration (*R*_g_) of 7.2 nm ± 0.1 nm (Fig. 1c and Table 1). Consistent with these values, the right-skewed pair distance distribution function, *P(r)*, reflects the extended nature of the molecule (Fig. 1c). To derive the low-resolution three-dimensional molecular envelope of G3BP1, we performed *ab initio* modeling from the SEC-SAXS data. P2 symmetry was applied throughout all calculations, to reflect the dimeric nature of G3BP1 in solution. The resulting *ab initio* model of G3BP1 revealed an elongated dimer characterized by a bulky center and distinct lateral extensions (Fig. 1d, upper inset). The *ab initio* model showed an excellent fit to the experimental data, with a *χ*^2^ of 0.99 (Fig. 1d, black line).

**Table 1.**
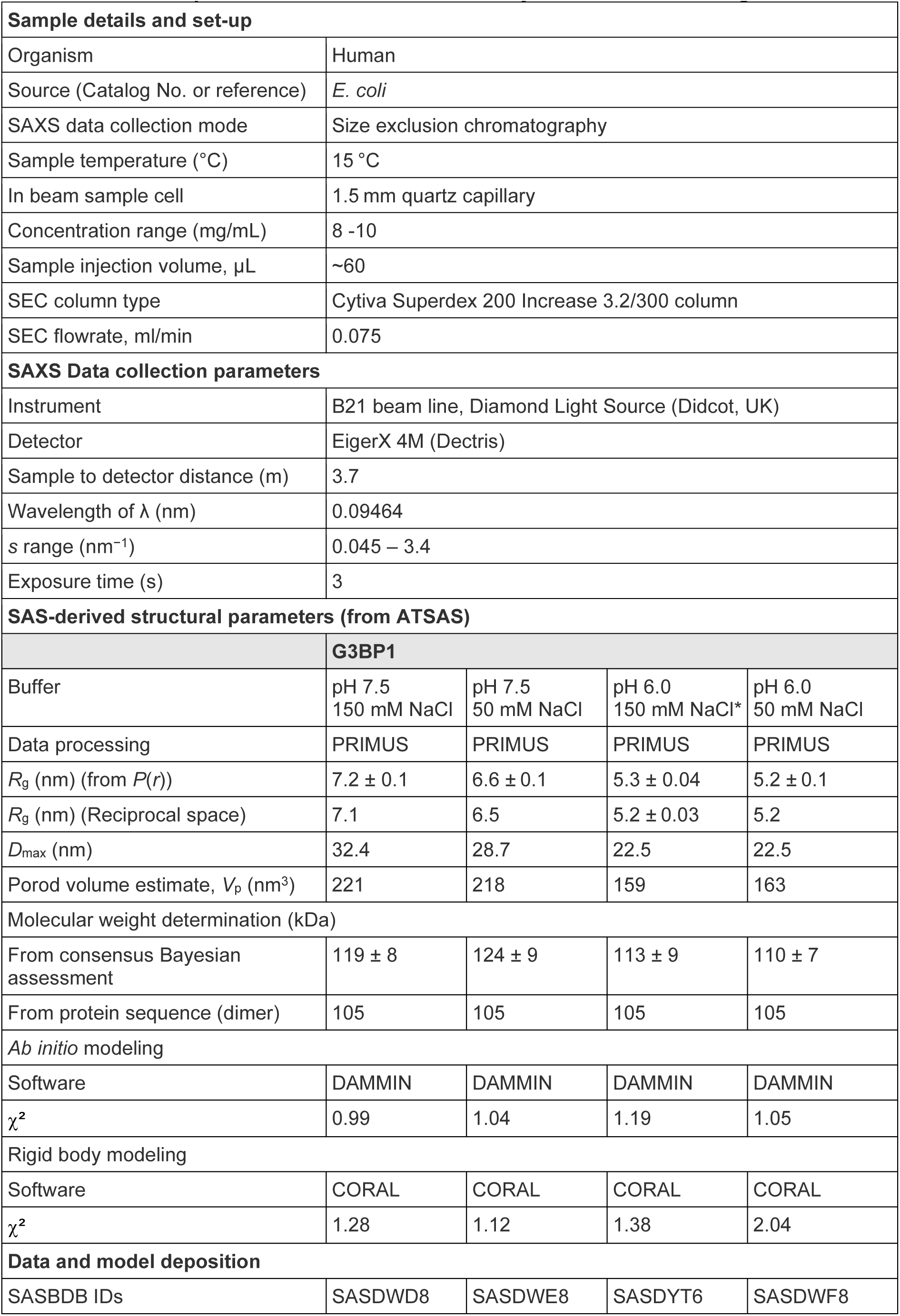
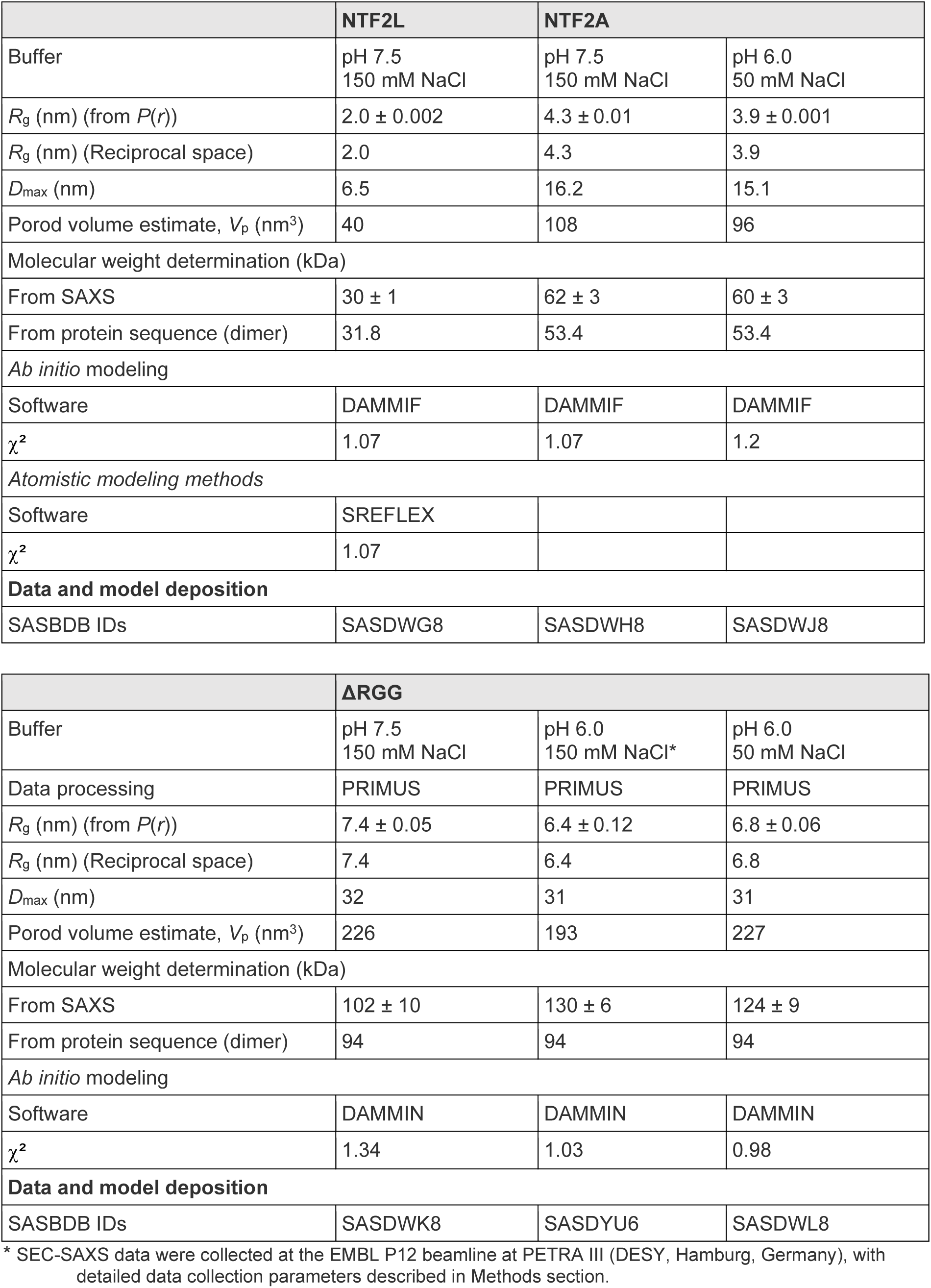
SAXS sample details, data collection, analysis, and 3D modeling details.

To further characterize the molecular architecture of the G3BP1 dimer, we performed SAXS-based hybrid rigid body modeling using the program CORAL. The crystal structure of the NTF2L homodimer (PDB ID: 6TA7)^39^ and a molecular model of the globular RRM (predicted using AlphaFold) were used as rigid bodies, while the remaining segments were represented as chains of dummy residues. A representative model is shown in Fig. 1d (lower inset), with a *χ*^2^ value of 1.28, which indicates a good fit to the scattering profile. Importantly, the hybrid model agrees well with the *ab initio* SAXS envelope of G3BP1 (Fig. 1d, upper inset) and suggests a global topology wherein the NTF2L dimer forms a central core, with the IDR1, PxxP, RRM, and RGG regions extending outward in opposite directions (Fig. 1d, lower inset). Notably, the IDRs do not appear as entirely random coils but instead adopt relatively constrained conformations.

To validate the domain arrangement of G3BP1 within the hybrid model, we purified the NTF2L domain. Additionally, we constructed a truncate comprising NTF2L and the adjacent IDR1 region, henceforth referred to as NTF2LA (Fig. 2a). SEC-SAXS data were subsequently collected for both NTF2L and NTF2LA to assess their individual structural contributions to the full-length assembly. The Kratky plot analyses indicated that NTF2L adopts a globular conformation, while NTF2LA exhibits a characteristically elongated conformation (Fig. 2b). Consistent with the enhanced flexibility conferred by IDR1, *P(r)* analysis showed an increase in *D*_max_, expanding from 7 nm (NTF2L) to 16 nm (NTF2LA) (Fig. 2c and Table 1). *Ab initio* models were subsequently calculated for both constructs (Fig. 2d). The NTF2L model adopted a globular shape that closely matched the NTF2L dimer crystal structure (Fig. 2d, light gray). In contrast, the *ab initio* model of NTF2LA displayed an elongated shape featuring a bulkier center with comparable size to the NTF2L dimer, whose morphology was consistent with the IDR1 region being positioned at each terminus (Fig. 2d, dark gray and Appendix Fig. S2). These results rule out the possibility of a full-length parallel G3BP1 dimer assembly, as such a configuration would require a significantly thicker and shorter shape than our experimental observations. Crucially, the available volume at the distal ends of the *ab initio* G3BP1 envelope is insufficient to accommodate the NTF2L dimer in a parallel orientation. The SAXS data and the structural models have been deposited into the Small Angle Scattering Biological Data Bank (SASBDB^40^) and the statistics are summarized in Table 1.

**Fig. 2:**
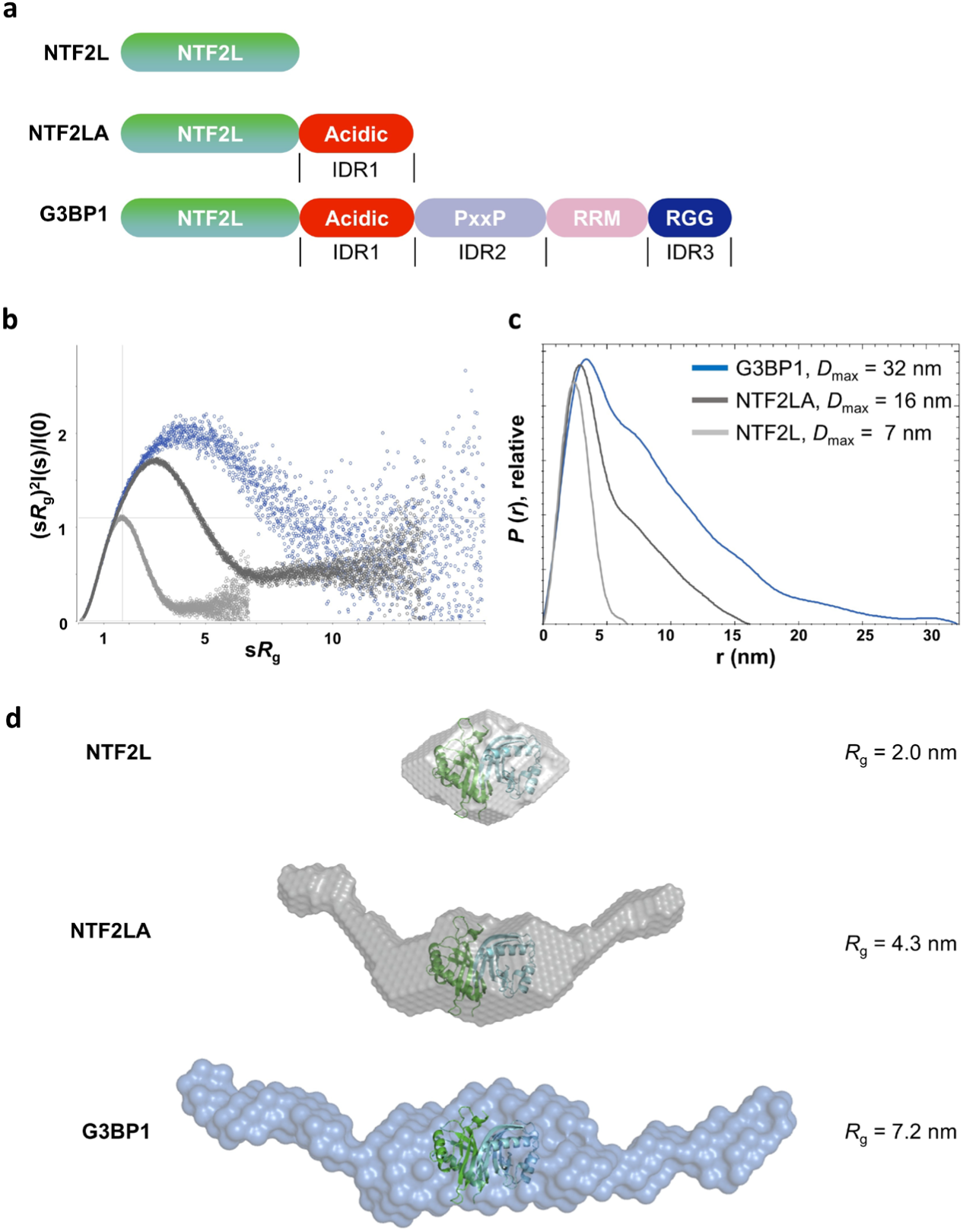
Comparative SEC-SAXS analyses of full-length and truncated G3BP1 variants confirm that G3BP1 forms a head-to-head antiparallel dimer. **a**, Schematic diagrams illustrating the domain organizations of NTF2L, NTF2LA and full-length G3BP1. **b**, Scattering curves of the three samples shown as normalized Kratky plots. **c**, Distance probability function, *P(r)* of the three samples, reveals distinct shapes and sizes for each construct. The curve of NTF2L is shown in light gray, NTF2LA in dark gray and full-length G3BP1 in blue. **d**, Overlay of the NTF2L dimeric crystal structure (PDB code 6TA7) onto the *ab initio* reconstructions. *Ab initio* models (generated with DAMMIF) are presented as spheres. SAXS data were collected in a buffer containing 20 mM Tris-HCl (pH 7.5) and 150 mM NaCl.

To summarize, the comprehensive analysis of the experimental SEC-SAXS data, combined with the truncates analysis, demonstrates that G3BP1 forms an antiparallel dimer in a head-to-head configuration, with the NTF2L homodimer centrally positioned within the overall structure (Fig. 1d). Crucially, the flexible IDRs are structurally constrained near the NTF2L domain and the RRM, resulting in an overall compact and well-defined topology for the full-length G3BP1 dimer.

### pH 6.0 drives G3BP1 conformational compaction

Previous studies have suggested that the conformation of G3BP1 is sensitive to pH and salt concentration^22,23^. In line with these observations, our recent work has demonstrated that acidification occurs within both cellular and reconstituted condensates^39^. Specifically, we observed a pH drop of at least 0.5-1.0 units within mammalian SGs compared to the adjacent cytoplasm. To systematically probe the microenvironmental influence on conformational dynamics of G3BP1, SEC-SAXS experiments were performed under different pH and ionic conditions: pH 7.5 with 150 mM NaCl, pH 7.5 with 50 mM NaCl, pH 6.0 with 150 mM NaCl and pH 6.0 with 50 mM NaCl (Fig. 3a).

**Fig. 3:**
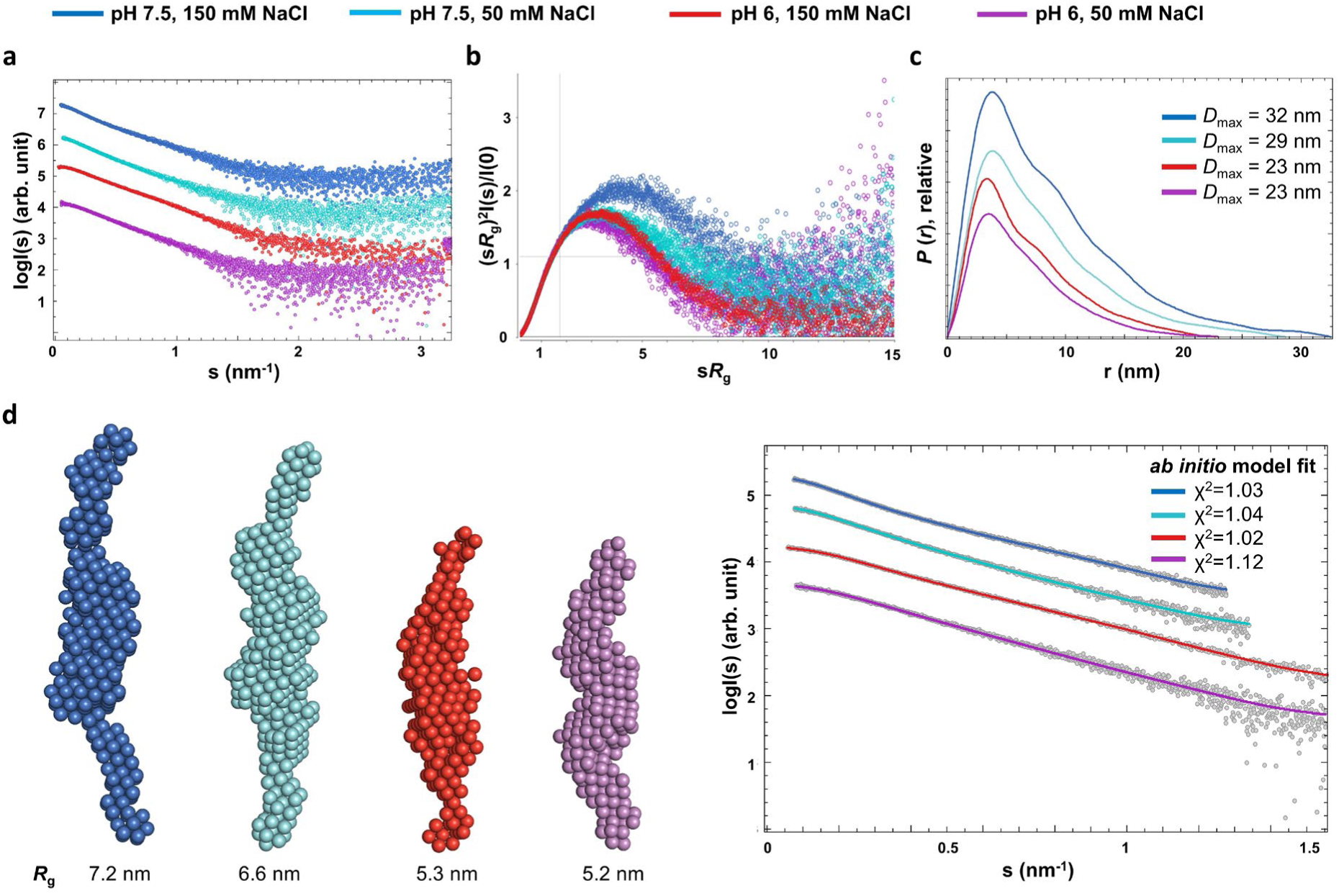
Full-length G3BP1 undergoes conformational compaction in response to pH drops. **a**, Scattering profiles of G3BP1 under indicated buffer conditions. Curves have been vertically shifted for better visibility. **b**, Representation of the scattering curves as Normalized Kratky plots. **c**, Distance distribution functions of the G3BP1 samples. **d**, *Ab initio* models (generated with DAMMIN) are presented as spheres in the left panel. The right panel shows the fits of the *ab initio* models to the experimental scattering data within the restricted angular range. Scattering profiles are represented as gray dots and the fitting of the *ab initio* models as colored lines.

The analysis of normalized Kratky plots displayed a leftward peak shift as the pH decreased from 7.5 to 6.0 (Fig. 3b), indicating a reduction in conformational flexibility. This trend was further corroborated by distance distribution functions, which showed a marked contraction in *D*_max_ from 32 nm (pH 7.5) to 23 nm (pH 6.0) (150 mM NaCl; Fig. 3c and Table 1). Consistent with this compaction, the *R*_g_ decreased from 7.2 ± 0.1 nm (pH 7.5) to 5.4 ± 0.1 nm (pH 6.0) (Table 1). Furthermore, the *ab initio* envelopes reconstructed from the scattering data at indicated conditions consistently illustrated that G3BP1 undergoes a pH-induced transition from an extended to a compact conformation (Fig. 3d). Moreover, since a reduction in salt concentration had no significant impact on the overall conformation at either pH, our findings underscore the dominant role of pH in driving this specific conformational transition.

### G3BP1 undergoes homotypic, RNA-independent phase separation at pH 6

Prompted by the pH-induced conformational compaction revealed by SAXS, we next investigated its functional impact on G3BP1 phase separation. Fluorescence microscopy was utilized to monitor the intrinsic phase separation of G3BP1 across a range of pH values in the absence of RNA. These *in vitro* LLPS assays were performed at a protein concentration of 5 µM, mimicking the physiological abundance of G3BP1 in mammalian cells^22,24^. Intriguingly, G3BP1 exhibited homotypic, RNA-independent phase separation at pH 6.0 (Fig. 4a). The observed droplets followed a distinct salt-dependent trend; phase separation was most pronounced at 50 mM NaCl, attenuated at 150 mM NaCl, and abolished at 300 mM NaCl.

**Fig. 4:**
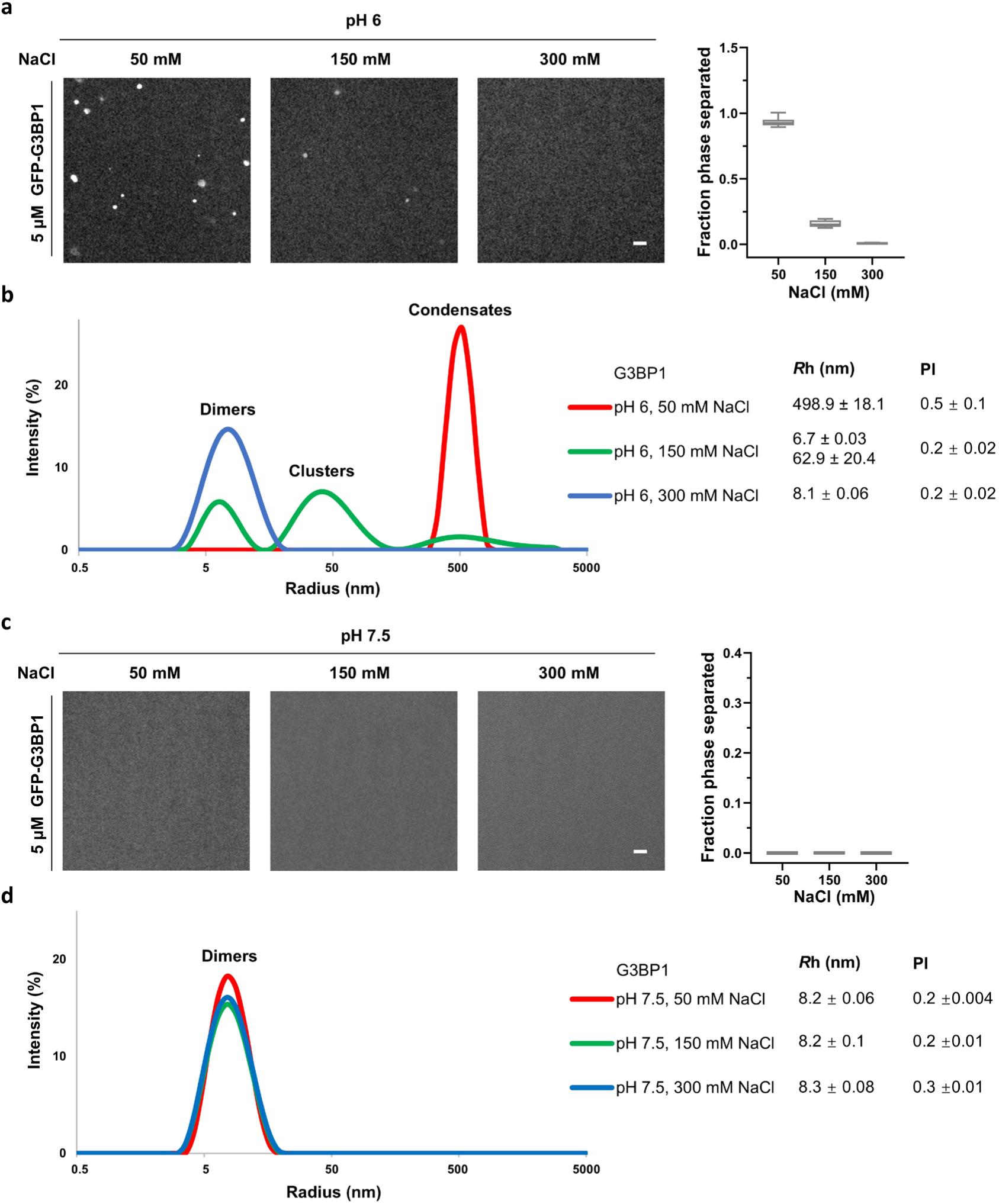
G3BP1 undergoes homotypic phase separation at pH 6.0. **a**, Representative fluorescence images of G3BP1 (5 µM) at pH 6.0, under indicated salt conditions. Quantification of phase-separated fraction is presented as mean ± SD (n = 10 fields of view, FOVs). Scale bar, 10 µm. **b**, DLS measurements of G3BP1 (5 µM) at pH 6.0 across indicated salt concentrations, showing the particle size distribution (mean by intensity) profiles. **c**, Representative fluorescence images of G3BP1 (5 µM) at pH 7.5, under indicated salt conditions. Quantification of phase-separated fraction is presented as mean ± SD (n = 10 FOVs). Scale bar, 10 µm. **d**, DLS measurements of G3BP1 (5 µM) at pH 7.5, under indicated salt concentrations, showing the particle size distribution (mean by intensity) profiles. Three independent DLS measurements were performed. Data represent the mean ± SD of three independent experiments.

DLS measurements further corroborated these observations by capturing the higher-order assembly of G3BP1 (Fig. 4b). Specifically, at pH 6.0 and 50 mM NaCl, the G3BP1 solution displayed a peak centered at approximately 500 nm, consistent with the formation of large homotypic condensates (Fig. 4b, red curve and Appendix Fig. S3a). At 150 mM NaCl, the equilibrium shifted toward smaller species, with the overall population mainly consisting of G3BP1 dimers and clusters (*R_h_* ≈ 62.9 nm) (Fig. 4b, green curve and Appendix Fig. S3b). At 300 mM NaCl, G3BP1 existed exclusively as monodisperse particles with a *R_h_* of ∼ 8.1 nm, matching the dimeric dimension at pH 7.5 (Fig. 4b, blue curve and Appendix Fig. S3c). In contrast, no homotypic phase separation was observed at pH 7.5 across the same salt range (Fig. 4c), where G3BP1 remained exclusively dimeric as verified by DLS (Fig. 4d).

A previous study interpreted the increased *Rh* at pH 6.0 as a sign of conformational expansion^22^. However, since DLS captures the total scattering signal from all species in solution, this larger *Rh* may instead reflect a change in the oligomerization state. Given our observation that G3BP1 undergoes homotypic phase separation at pH 6.0, this increase likely reflects the formation of higher-order clusters or condensates, rather than a genuine conformational expansion of the G3BP1 dimer.

### pH 6 enhances RNA-mediated phase separation of G3BP1

Given that SG assembly is closely correlated with a sudden increase in free cytosolic RNA^22,23,29^, we next examined the phase separation behavior of G3BP1 in the presence of RNA to better reflect the cellular environment during stress. At 7.5, the G3BP1-RNA condensates were sensitive to ionic strength. While condensates were clearly observed at pH 7.5 and 50 mM NaCl, they were absent at physiological (150 mM) and high (300 mM) salt concentrations (Fig. 5a), which aligns with previous reports^22,23^. These results were confirmed using untagged G3BP1, to exclude any potential interference from the GFP tag (Appendix Fig. S4a). Furthermore, we monitored the real-time dynamics of this process by titrating NaCl into pre-formed condensates. The rapid disassembly of G3BP1-RNA condensates upon increasing the NaCl concentration to 150 mM (Fig. 5b) suggests that G3BP1-RNA condensation at physiological pH can be reversibly modulated by salt concentration.

**Fig. 5:**
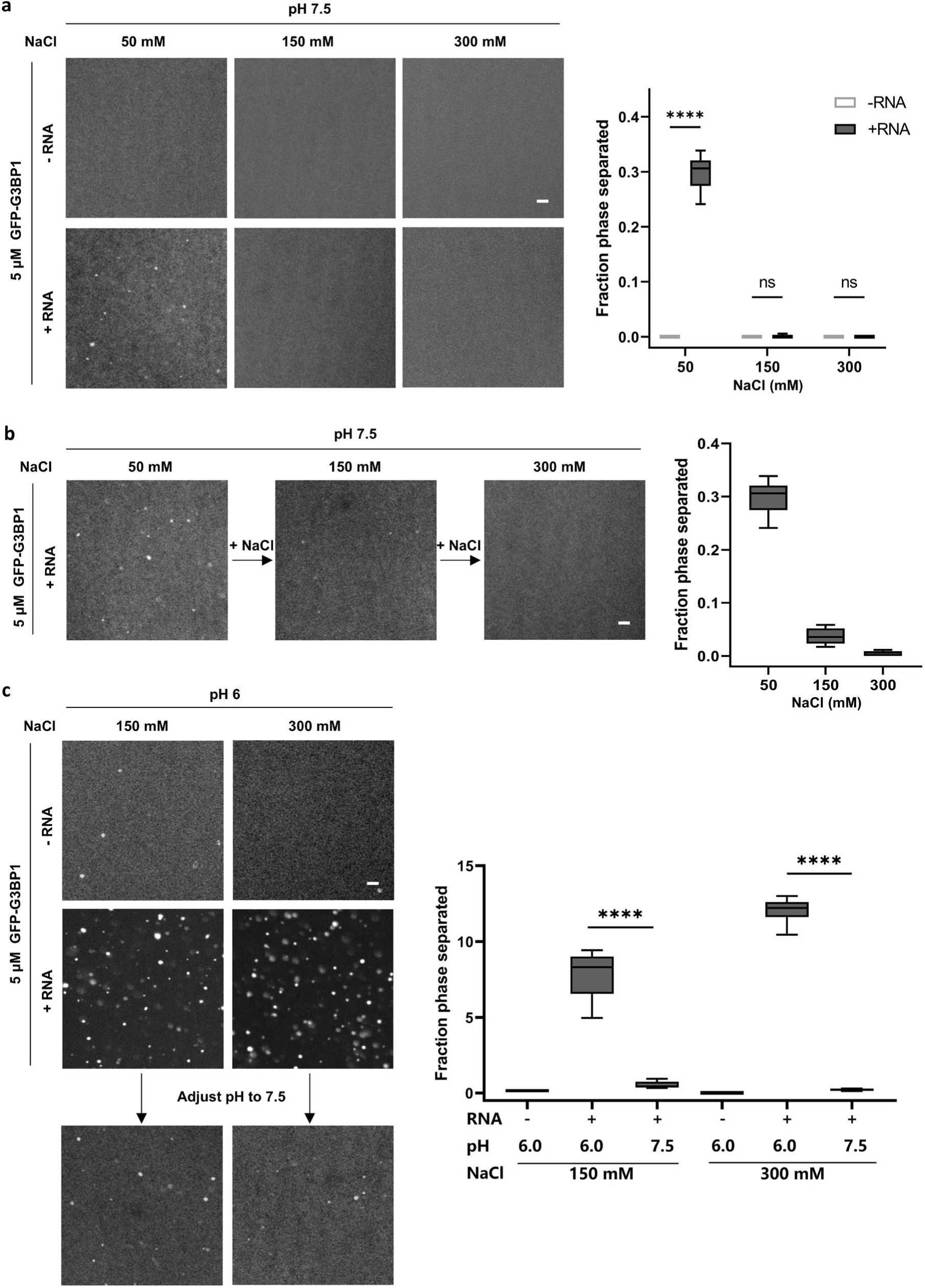
pH 6.0 enhances RNA-mediated phase separation of G3BP1. **a**, Representative fluorescence images of G3BP1 condensates with or without total RNA (50 ng/µL) at pH 7.5, under indicated salt conditions. Quantification of phase-separated fraction is presented as mean ± SD (n = 10 FOVs). Scale bar, 10 µm. **b**, Representative fluorescence images and quantification of G3BP1-RNA condensates during salt titration at pH 6.0. **c**, Representative fluorescence images of G3BP1 condensates with or without total RNA (50 ng/µL) at pH 6.0, under indicated salt conditions or during pH titration. Quantification of phase-separated fraction is presented as mean ± SD (n = 10 FOVs). Scale bar, 10 µm.

Conversely, acidic conditions significantly enhanced the RNA-mediated LLPS of G3BP1 (Fig. 5c). While G3BP1-RNA condensates were absent at pH 7.5 and 150 mM NaCl, they formed with high efficiency at pH 6.0 (150 mM NaCl; Fig. 5a). These acidic condensates, notably, exhibited remarkable salt resistance, persisting even at 300 mM NaCl (Fig. 5c). Similar results were obtained using untagged G3BP1 (Appendix Fig. S4b). This resilience to high ionic strength suggests that the pH-induced compact conformation promotes interaction networks beyond simple electrostatics, likely involving hydrophobic effects, π-π stacking, and cation-π interactions, forces often mediated by IDRs^41,42^. In addition, G3BP1-RNA condensates formed under acidic conditions rapidly dissolved when the pH was titrated to 7.5 (Fig. 5c), demonstrating the reversibility of this pH-modulated assembly.

The physiological relevance of acidification-enhanced G3BP1 condensation is evident within the context of endolysosomal damage. Following lysosomal leakage, a sudden localized decrease in cytosolic pH (from 7.5 to below 6.5) promotes the rapid formation of G3BP1 condensates, which are essential for stabilizing ruptured membranes and facilitating the efficient repair of damaged endolysosomes^43^. The pH-dependent conformational switch of G3BP1 likely serves as a biophysical regulator, increasing its effective valency in response to acidification. Such a mechanism would ensure the rapid, site-specific assembly of G3BP1 at stress loci where localized pH fluctuations and the release of untranslated mRNA coincide.

Overall, these findings demonstrate that pH 6.0 drives a distinct conformational compaction of G3BP1, a transition that coincides with the emergence of homotypic phase separation and the robust enhancement of RNA-mediated LLPS. This pH-responsive structural plasticity provides a compelling biophysical mechanism, explaining how G3BP1 assemblies are dynamically tuned by pH fluctuations during the cellular stress response.

### The RGG region is essential for acidity-driven G3BP1 conformational compaction

Previous studies have indicated that interactions between IDRs are crucial for regulating G3BP1-centered SG assembly^22,23^. The deletion of the intrinsically disordered RGG region prevents SG formation^21^, despite the truncated variant retaining its RNA-binding capacity^22^. To investigate the contribution of the RGG region to the conformational and functional dynamics of G3BP1, we purified a truncated variant lacking this region, termed ΔRGG (Fig. 6a), and performed SEC-SAXS analysis under the same conditions as for the full-length G3BP1 (Fig. 6b-g). At pH 7.5, ΔRGG adopted an extended conformation identical to full-length G3BP1, characterized by comparable *D*_max_ and *R*_g_ values (Fig. 6 and Table 1). Furthermore, the *ab initio* models confirmed that ΔRGG maintained an elongated shape as the full-length protein at neutral pH (Fig. 6e, i and iii). Crucially, unlike the full-length protein, ΔRGG failed to undergo conformational compaction at pH 6.0 (Fig. 6e, ii and iv), with its *D*_max_ remaining largely unchanged (Fig. 6d).

**Fig. 6:**
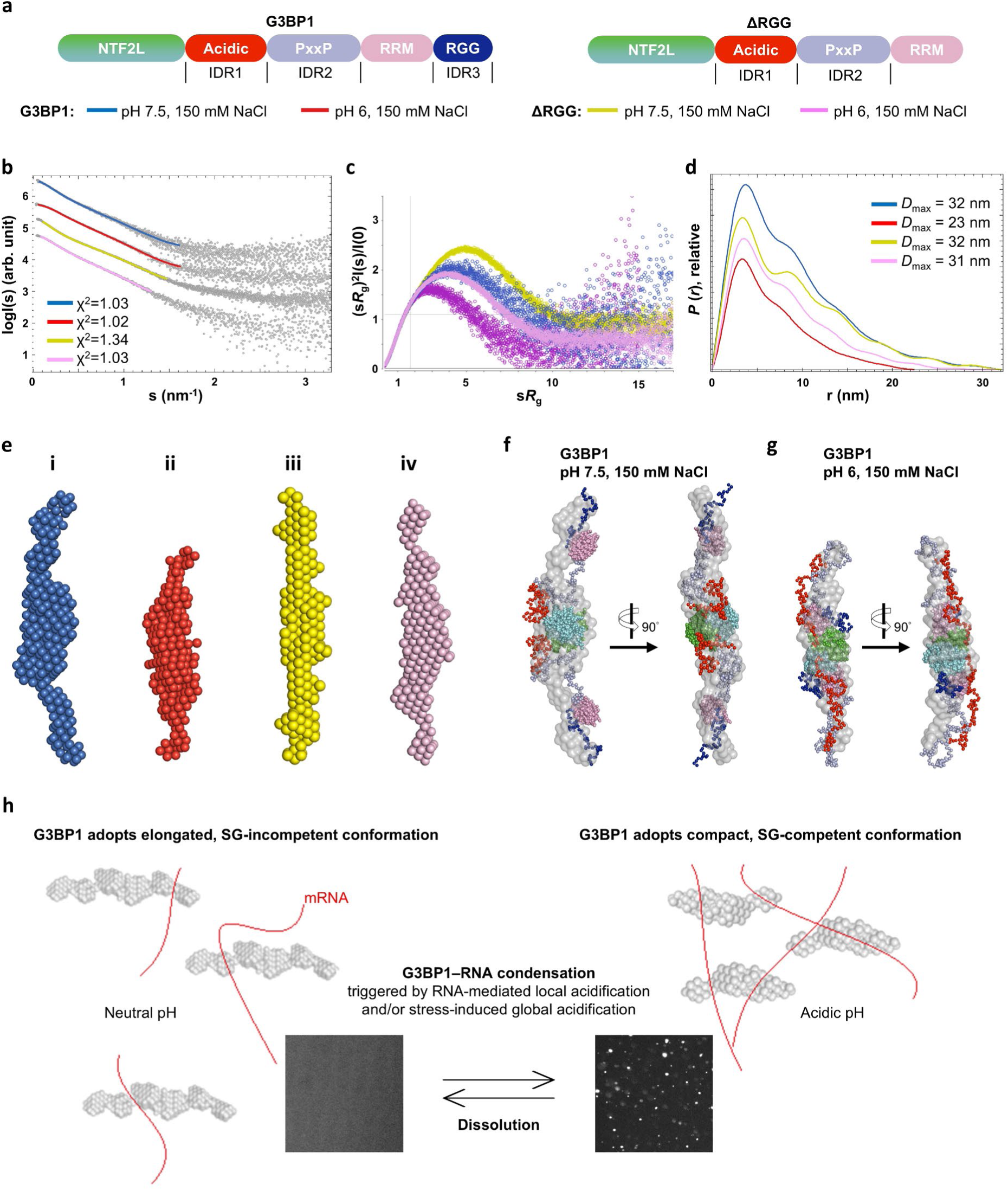
The RGG region is essential for acidity-induced conformational compaction of G3BP1. **a**, Schematic diagrams illustrating the domain organization of G3BP1 and the RGG-deleted mutant (ΔRGG). **b**, Scattering curves and the fits of the *ab initio* models to the experimental scattering data for G3BP1 (i and ii) and ΔRGG (iii and iv) under indicated conditions. Scattering profiles are represented as gray dots and the fitting of the *ab initio* models as colored lines. Curves have been vertically shifted for better visibility. **c**, Scattering curves are shown in normalized Kratky plot. **d**, Distance distribution functions of G3BP1 and ΔRGG. **e**, *Ab initio* models (generated with DAMMIN) of G3BP1 and ΔRGG under indicated conditions are presented as spheres. The fits of the *ab initio* models to the experimental scattering data within the restricted angular range are shown in (**b**). **f**, Superimposition of the *ab inito* and hybrid models of G3BP1 obtained using DAMMIN and CORAL, respectively. The *ab inito* models are represented as gray semitransparent spheres, hybrid models are shown as spheres, with the domains colored as indicated in (**a**). **g**, A mechanistic model linking cytoplasmic pH drop and mRNA availability to SG formation via the G3BP1 conformational switch.

To provide a structural basis for G3BP1 conformational dynamics, we performed CORAL hybrid modeling integrated with experimental SAXS data collected at pH 7.5 and 6.0. For full-length G3BP1, independent modeling runs yielded models with similar spatial arrangements that fit the experimental scattering curves (Appendix Fig. S6) and were in close agreement with the *ab initio* envelopes (Fig. 6f, g). A comparison of these hybrid models suggests that pH reduction triggers a pronounced spatial rearrangement of G3BP1’s IDRs. Specifically, while IDR1 appears to be more extended, the RRM and RGG region move closer to NTF2L central core at pH 6.0, resulting in a significantly more compact and shortened global conformation (Fig. 6f, g and Appendix Fig. S6b, c). This observation aligns with a previous *in silico* study suggesting that the G3BP1 compaction correlates linearly with the inter-domain spacing between IDR1 and the RGG region^23^. Collectively, these results indicate that the RGG region is indispensable for G3BP1’s conformational compaction in response to acidification.

The functional consequence of this structural defect was confirmed via systematic LLPS assays using tag-free ΔRGG. No droplet formation was observed at pH 7.5 regardless of the presence of RNA (Appendix Fig. S5a), as is consistent with previous reports^21,23^. Notably, at pH 6.0, ΔRGG failed to undergo homotypic condensation, and its RNA-mediated condensation was profoundly attenuated compared to the full-length protein (Appendix Fig. S5b and Figs, 4a, 5c). DLS analysis revealed that ΔRGG remains dimeric at pH 7.5, mirroring the behavior of the full-length protein (Appendix Fig. S5c). However, ΔRGG lost the capacity to assemble into large-scale condensates at pH 6, forming only small intermediate clusters at low salt concentrations (Appendix Fig. S5d).

Altogether, these findings reveal that G3BP1 phase behavior is intrinsically coupled to its conformational state. We show that RGG-dependent conformational compaction acts as a structural prerequisite that primes G3BP1 for efficient homotypic and RNA-dependent condensation. By undergoing this pH-triggered ’expanded-to-compact’ transition, G3BP1 amplifies its effective valency for both self-association and protein-RNA interactions, thereby facilitating rapid and site-specific condensate formation in response to localized acidification.

### A biophysical model: Acidification triggers G3BP1 conformational compaction to seed initial condensate assembly

Previous studies have established that various stressors, such as hypoxia, heat shock and nutrient deprivation, can trigger intracellular pH drop in mammalian, plant, and microbial cells^44–48^. In response to stress-induced pH changes, specific proteins and RNAs undergo LLPS to promote cellular survival and adaptation. Additionally, intracellular acidification is intimately linked to aging processes and aging-related neurodegenerative diseases^49–51^.

Building upon our previous experimental evidence that cellular SGs are indeed more acidic than the surrounding cytoplasm^39^ and with the findings of the current study, we propose a biophysical model in which a pH-triggered conformational switch in G3BP1 dictates its initial phase separation capacity (Fig. 6h). Under physiological conditions at neutral pH, G3BP1 adopts an extended dimeric conformation which inherently disfavors spontaneous condensation, thereby maintaining a diffuse cytoplasmic distribution. However, during cellular stress, translational arrest leads to the accumulation of free mRNA. The high density of these polyanionic RNA molecules creates a strong negative electrostatic potential that locally sequesters counterions^52,53^, primarily protons (H^+^), consequently establishing a pH drop in the immediate vicinity of G3BP1. The resulting acidic microenvironment drives the conformational compaction of G3BP1, amplifying its effective valency for both homotypic and RNA-mediated heterotypic LLPS. By coupling localized mRNA cues with local pH fluctuations, this integrated mechanism ensures the rapid and site-specific initiation of G3BP1 condensation, providing a robust and tunable biophysical framework for cellular stress adaptation.

## Discussion

The dynamic assembly and disassembly of RNP granules are fundamental to maintaining cellular homeostasis, particularly under stress conditions. Dysregulation of these granules, including alterations in their structural integrity and turnover dynamics, has been implicated in tumor progression and other severe human diseases^3–5^. As the central node and the molecular switch initiating SG formation, G3BP1 is functionally indispensable. However, a precise mechanistic understanding of SG assembly has long been impeded by the lack of structural information on full-length G3BP1. The extensive IDRs and inherent conformational heterogeneity of the full-length protein preclude characterization by traditional X-ray crystallography or cryo-electron microscopy. Consequently, current knowledge relies heavily on hypothetical models derived from truncated domains or computational predictions.

Our results address this structural ambiguity and reveal that full-length G3BP1 adopts an elongated, head-to-head antiparallel homodimer architecture in solution (Figs. 1 and 2). Crucially, we demonstrate that this architecture is not static but functions as a pH-responsive sensor, where acidic environments trigger an RGG-dependent conformational compaction (Figs. 3 and 6). This transition into a compact LLPS-competent state provides a direct mechanistic link between cytoplasmic acidification and the physical initiation of condensate assembly (Figs. 4, 5 and Appendix Fig. S5). In this context, the RGG domain not only acts as an auxiliary RNA-binding element but also a structural regulator of the conformational plasticity of G3BP1, thereby tuning the biophysical threshold for phase separation.

SG assembly correlates with the rapid release of mRNAs from polysomes following translational arrest^26,27^. Our findings suggest a direct physicochemical coupling between this increase of free mRNA and the activation of G3BP1. We propose that the polyanionic nature of accumulated RNA generates a local acidic microenvironment, inducing a conformational compaction in G3BP1 that enables rapid phase separation. Subsequent clearance of free RNA and restoration of physiological pH would favor the reversion to the elongated conformation of G3BP1, thereby attenuating multivalent interactions and promoting condensate dissolution.

In conclusion, our study provides the first comprehensive structural characterization of full-length G3BP1, establishing an experimentally validated framework for its conformational dynamics. Crucially, we reveal that acidic environments drive a reversible conformational compaction of G3BP1 that promotes phase separation. This mechanism likely operates in synergy with cytoplasmic acidification, a hallmark of various physiological stresses^44–48^ and aging-related neurodegenerative diseases^49–51^. By sensing these intracellular pH fluctuations, G3BP1 functions as a molecular sensor, establishing a direct biophysical link between conformational plasticity and the regulation of condensate homeostasis. Ultimately, by resolving the structural basis of G3BP1-mediated condensate assembly, our work paves the way for the rational design of allosteric inhibitors and opens promising therapeutic avenues for addressing pathologies driven by aberrant stress granule dynamics.

## Methods

### Experimental model and study participant details

#### Escherichia coli strains

Recombinant proteins were expressed in *Escherichia coli* BL21(T7 Express lysY/Iq) host cells, grown in terrific broth (TB) medium at 25°C, 220 rpm.

### Cloning, protein expression and purification

The DNA fragment encoding full-length human G3BP1 was PCR-amplified from a U2OS cDNA library and subsequently subcloned into the pET30 vector using ligation-independent cloning^54^. To facilitate purification, a TEV cleavage site was introduced between the N-terminal poly-His6 tag and G3BP1. The final construct, termed his6-TEV-G3BP1, was validated by DNA sequencing (Eurofins Genomics) and served as the template for constructing the G3BP1 deletion mutants used in this study. The expression constructs of his6-TEV-NTF2L (residues 1-139), his6-TEV-NTF2LA (residues 1-232), his6-TEV-G3BP1ΔRGG (residues 1-413) and his6-EGFP-TEV-G3BP1 were created using the same cloning method and verified by DNA sequencing.

Full-length recombinant G3BP1 was expressed in T7 Express lysY/Iq competent *E. coli* cells at 25°C overnight in terrific broth (TB) and purified under native conditions. Bacteria cells were grown to an OD of approximately 0.7 in TB medium and induced with 0.5 mM IPTG at 25 °C overnight, then pelleted and resuspended in lysis buffer containing 50 mM HEPES (pH 7.5), 300 mM NaCl, 1 mM DTT, complete protease inhibitor cocktail tablet (Roche), and Benzonase Nuclease (Millipore). Cells were lysed by sonication (10 minutes, 40% amplitude, 5 seconds on, 5 seconds off). After centrifugation to remove cell debris, the resulting supernatant was applied to Ni-NTA agarose resin (QIAGEN) for affinity purification. The eluent was collected, and the poly-His tag was removed by TEV protease (PSF, KI, Stockholm) in HEPES buffer containing 0.5 mM EDTA and 1 mM DTT. The cleaved protein was passed through the Ni-NTA column as flow-through. Subsequently, G3BP1 was loaded onto a HiTrap Heparin HP affinity column (Cytiva) to eliminate non-specific RNA binding. The fractions were analyzed by SDS-PAGE, and the corresponding tubes containing G3BP1 were pooled and concentrated. Benzonase Nuclease was included during the purification process, and the OD260/280 ratio of batch purified G3BP1 ranged from 0.58 to 0.6, indicating efficient removal of nucleic acid contaminants. Full-length G3BP1 was purified using a Superdex 200 increase10/300 GL column (Cytiva) in the desired buffers for SAXS studies. The monodisperse G3BP1 fractions were concentrated, filtered, flash frozen in liquid nitrogen, and stored at −80°C. His6-TEV-ΔRGG (residues 1-413) and his6-EGFP-TEV-G3BP1 were purified using similar protocols. To assess the degree of polydispersity of the freshly purified proteins, DLS was performed and the hydrodynamic diameter of the proteins was determined.

Deletion constructs NTF2L and NTF2LA were purified as previously described.^35^ Briefly, the NTF2L dimer was affinity-purified using Ni-NTA agarose resin and isolated via Superdex 75 10/300 GL size exclusion chromatography (GE Healthcare). Following TEV cleavage, NTF2L was collected as the Ni-NTA flow-through. Both NTF2L and NTF2LA were further purified using a Superdex 75 10/300 GL column after tag removal to enhance purity. Protein-containing fractions were pooled and concentrated using 10 kDa MWCO Amicon ultra centrifugal filters (Millipore).

### Small Angle X-ray Scattering (SAXS)

SAXS data were recorded at the B21 beamline of the Diamond Light Source (Didcot, United Kingdom)^55^. Data collection occurred at 3-second intervals using an EigerX 4M detector (Dectris, Switzerland) at a sample detector distance of 3.7 m and at a wavelength of λ = 0.09464 nm (I(s) vs s, where s = 4π*sinθ/λ, and 2θ is the scattering angle), with fixed energy at 13.1 keV, covering the scattering vector range from 0.045 to 3.4 nm⁻¹. For the SEC-SAXS runs of G3BP1, an Agilent 1200 HPLC system equipped with a Superdex 200 increase 3.2/300 column (Cytiva) was used. 60 µL of the sample was injected and run at flow rate of 0.075 ml/min in indicated buffer. The SEC-separated sample was then exposed to X-rays in a 1.5-mm diameter, 10-μm-thick quartz capillary flow cell. The running buffer was supplemented with 1% glycerol to mitigate radiation damage, samples and the corresponding matched solvent blank were measured.

Additional SEC-SAXS data of G3BP1 and ΔRGG in buffer pH 6, 150 mM NaCl were collected on the EMBL P12 beamline at PETRA III (DESY, Hamburg, Germany^56^) using a Pilatus 6 M detector at a sample-detector distance of 3 m and at a wavelength of *λ* = 0.124 nm (I(s) vs s, where s = 4πsinθ/λ, and 2θ is the scattering angle). The SEC parameters were as follows: 60 μl sample at indicated concentration (Table 1) was injected at a flow rate of 0.35 ml/min onto a GE Superdex 200 Increase 5/150 column at 20 °C. 1800 successive frames, each with 0.5 s exposure times, were collected. The running buffer was supplemented with 1% glycerol to reduce radiation damage, samples and the corresponding matched solvent blank were measured.

### SAXS data processing and modeling

Initial SEC-SAXS datasets were processed using Chromix^57^, both automatically and interactively, resulting in buffer subtracted curves. Further data processing was then performed with the ATSAS 3.1 suite^58^. The radius of gyration (R*_g_*) was obtained by fitting the linear Guinier region of the data using PRIMUS^59^. The pair distribution function, *P*(*r*), was generated with GNOM^60^. *Ab initio* models of molecular envelopes were generated using DAMMIN^61^ or DAMMIF^62^, where scattering from the calculated envelopes was fitted against the experimental curves and evaluated by the χ² values. The resulting twenty *ab initio* models were averaged with DAMAVER.^63^ Advance hybrid modeling using available high resolution parts of the structure with addition of the missing loops/domains was conducted with CORAL^64^. A SASpy plugin in PyMOL^65^ was employed to overlay the dimeric NTF2L X-ray structure with the corresponding *ab initio* models. The SAXS data are summarized in Table 1 and the respective curves and models have been deposited into SASBDB^40^.

### *In vitro* liquid–liquid phase separation (LLPS) assays

For G3BP1-RNA condensate formation, 5 μM of EGFP-G3BP1 or 25 μM of untagged protein sample were incubated with or without purified total RNA at varied buffer conditions. Total RNA was isolated from U2OS cells by TRIzol® reagent (Invitrogen, USA) and quantified with a Thermo Scientific NanoDrop 2000 spectrophotometer. Typically, 50 ng/µL total RNA was added to induce LLPS of G3BP1. The samples were mixed in low binding tubes (COSTAR 3206) and transferred to a 384-well ultra-low attachment microplate (Greiner Bio-one, Germany) at room temperature. Phase separation was visualized using a Nikon CrEST X-Light V3 Confocal Microscope equipped with a 20× air objective, controlled by NIS-Elements software. For EGFP-G3BP1 condensates, the spinning disk configuration with 488 nm excitation was employed, with a region of interest (ROI) of 1024 x 1024 pixels to facilitate rapid imaging and minimize photobleaching. High-resolution transmitted-light images of untagged G3BP1 condensates were acquired using a TL camera, focusing on a ROI measuring 610 x 640 pixels. Condensates were imaged with identical settings between replicates to allow for comparison of granule intensities. Images were captured within 10 minutes after mixing the protein, RNA and the indicated buffer, acquired as Z-stacks with a step size of 6–8 µm over a total depth of 200 µm (25–30 Z-planes per stack). Maximum intensity projections of the Z-stack images and subsequent quantitative analyses were performed using semi-automated workflows Fiji^66^. Scale bars were added through Analyze → Tools → Scale Bar function of Fiji. Three independent LLPS assays are performed.

### Dynamic light scattering (DLS)

Dynamic light scattering of protein samples was performed at 25°C with a Zetasizer Ultra instrument (Malvern Panalytical) using side scattering. Protein samples were mounted using 1.0 × 1.0 mm disposable cuvette capillaries (Malvern Panalytical) and placed within a low-volume disposable sizing cell kit holder (Malvern Panalytical). Typically, 5 µM protein samples were tested under the indicated pH and salt buffer conditions. Monodispersity and statistical size distribution were analyzed using the manufacturer supplied ZS Xplorer software. Mean ± SD data were calculated from three independent experiments.

### Quantification and statistical analysis

Statistical analyses of *in vitro* LLPS were performed using semi-automated workflows in Fiji. Particle size distribution analyses were performed using a Zetasizer Ultra and data were analyzed directly in the accompanying ZS Xplorer software. The statistical details for all experiments, including error bars, statistical significance, and n numbers, are provided in the figure legends and method details. Statistical comparisons between two groups were conducted using the unpaired t test (two tailed). Statistical analysis was conducted using GraphPad Prism 9.5.1 software. ns, P>0.05; *, P≤0.05; **, P≤0.01; ***, P≤0.001; ****, P≤0.0001.

The phase separation observed under each condition was quantified by analyzing 9-10 fields of view. Each 2048 x 2048 px frame was divided into 256 x 256 px tiles to account for any potential non-uniformity in condensation. Within each tile the fraction of phase separation was quantified as 𝐼_condensate_ · 𝐴_condensate_(𝐼_tile_ · 𝐴_tile_)^−1^and averaged to obtain a value for the whole frame. Condensation was systematically identified by applying a 9 x 9 px Gaussian blur to the tiles and filtering with an adaptive threshold of 5 plus the Gaussian-weighted sum the 21 x 21 px neighborhood around each pixel.

### Data and code availability

All relevant data associated with the published study are present in the paper or the Supplementary Information. The SAXS data generated in this study have been deposited in the SASBDB database (www.sasbdb.org) under accession codes <u>SASDWD8</u>, <u>SASDWE8</u>, <u>SASDWF8</u>, <u>SASDWG8</u>, <u>SASDWH8</u>, <u>SASDWJ8</u>, <u>SASDWK8</u>, <u>SASDWL8</u>, <u>SASDYT6</u> and <u>SASDYU6</u>.

## Acknowledgements

We acknowledge the Diamond Light Source for the allocation of beamtime (proposal mx29948-5) and for provision of synchrotron radiation facilities. The facility was supported by the Engineering and Physical Sciences Research Council (EPSRC) under grant number EP/R042683/1. We would like to thank the local coordinator Jodie Lavender and staff at the beamline Diamond B21 for assistance and support SAXS data collection. Part of the SAXS data were collected at beamline P12 operated by EMBL Hamburg at the PETRA III storage ring (DESY, Hamburg, Germany). We thank the beamline staff for their technical support. Part of this work was facilitated by the Protein Science Facility at Karolinska Institute, Stockholm, and we would like to thank Dr Emilia Strandback, Dr Henry Ampah-Korsah and Dr Tomas Nyman for assistance. This work was supported by grants from the Swedish Research Council (AA; 2021-05061), The Swedish Cancer Society (AA; 24 3775 Pj O1 H), The King Gustaf V Jubileum Fund (AA; 244092) and the Swedish Cancer and Allergy Foundation (AA; 11338).

## Author contributions

A.A., X.H., R.S. and D.S. conceptualized the project. X.H. and R.S. cloned, expressed, and purified all protein constructs. X.H. and R.S. prepared the SEC-SAXS samples, analyzed the scattering data and prepared the figures. M.A.G., C.E.B. and D.S. supervised and provided help with the analysis of the SEC-SAXS results. X.H. and R. S. conducted the LLPS assays in close collaboration with Q.Z. and M.F.; X.H., R. S., H.G.L., T.R. and A.A. wrote the manuscript. All authors discussed and commented on the final version. A.A., E.A., D.S., and H.G.L. have acquired all financial support.

## References

1. Alberti, S., Gladfelter, A. & Mittag, T. Considerations and Challenges in Studying Liquid-Liquid Phase Separation and Biomolecular Condensates. Cell 176, 419–434 (2019).

2. Posey, A.E. & Pappu, R.V. A First Glimpse of Nucleation of Phase Transitions in Living Cells. Mol Cell 71, 1–3 (2018).

3. Kedersha, N., Ivanov, P. & Anderson, P. Stress granules and cell signaling: more than just a passing phase? Trends Biochem Sci 38, 494–506 (2013).

4. Ripin, N. & Parker, R. Formation, function, and pathology of RNP granules. Cell 186, 4737–4756 (2023).

5. Zhou, H. et al. Stress granules: functions and mechanisms in cancer. Cell Biosci 13, 86 (2023).

6. Cui, Q., Liu, Z. & Bai, G. Friend or foe: The role of stress granule in neurodegenerative disease. Neuron 112, 2464–2485 (2024).

7. Hofmann, S., Kedersha, N., Anderson, P. & Ivanov, P. Molecular mechanisms of stress granule assembly and disassembly. Biochim Biophys Acta Mol Cell Res 1868, 118876 (2021).

8. Mateju, D. et al. Single-Molecule Imaging Reveals Translation of mRNAs Localized to Stress Granules. Cell 183, 1801–1812 e13 (2020).

9. Protter, D.S.W. & Parker, R. Principles and Properties of Stress Granules. Trends Cell Biol 26, 668–679 (2016).

10. Advani, V.M. & Ivanov, P. Stress granule subtypes: an emerging link to neurodegeneration. Cell Mol Life Sci 77, 4827–4845 (2020).

11. Gao, X. et al. Stress granule: A promising target for cancer treatment. Br J Pharmacol 176, 4421–4433 (2019).

12. McCormick, C. & Khaperskyy, D.A. Translation inhibition and stress granules in the antiviral immune response. Nat Rev Immunol 17, 647–660 (2017).

13. Brownsword, M.J. & Locker, N. A little less aggregation a little more replication: Viral manipulation of stress granules. Wiley Interdiscip Rev RNA 14, e1741 (2023).

14. Wolozin, B. & Ivanov, P. Stress granules and neurodegeneration. Nature Reviews Neuroscience 20, 649–666 (2019).

15. Farny, N.G., Kedersha, N.L. & Silver, P.A. Metazoan stress granule assembly is mediated by P-eIF2alpha-dependent and -independent mechanisms. RNA 15, 1814–21 (2009).

16. Franchini, D.M. et al. Microtubule-Driven Stress Granule Dynamics Regulate Inhibitory Immune Checkpoint Expression in T Cells. Cell Rep 26, 94–107 e7 (2019).

17. Garcia, M.A. et al. The chemotherapeutic drug 5-fluorouracil promotes PKR-mediated apoptosis in a p53-independent manner in colon and breast cancer cells. PLoS One 6, e23887 (2011).

18. Li, H. et al. MG53 suppresses tumor progression and stress granule formation by modulating G3BP2 activity in non-small cell lung cancer. Mol Cancer 20, 118 (2021).

19. Baron, D.M. et al. Amyotrophic lateral sclerosis-linked FUS/TLS alters stress granule assembly and dynamics. Mol Neurodegener 8, 30 (2013).

20. Sidibe, H., Dubinski, A. & Vande Velde, C. The multi-functional RNA-binding protein G3BP1 and its potential implication in neurodegenerative disease. J Neurochem 157, 944–962 (2021).

21. Kedersha, N. et al. G3BP-Caprin1-USP10 complexes mediate stress granule condensation and associate with 40S subunits. J Cell Biol 212, 845–60 (2016).

22. Guillen-Boixet, J. et al. RNA-Induced Conformational Switching and Clustering of G3BP Drive Stress Granule Assembly by Condensation. Cell 181, 346–361 e17 (2020).

23. Yang, P. et al. G3BP1 Is a Tunable Switch that Triggers Phase Separation to Assemble Stress Granules. Cell 181, 325–345 e28 (2020).

24. Sanders, D.W. et al. Competing Protein-RNA Interaction Networks Control Multiphase Intracellular Organization. Cell 181, 306–324 e28 (2020).

25. Tourriere, H. et al. The RasGAP-associated endoribonuclease G3BP assembles stress granules. J Cell Biol 160, 823–31 (2003).

26. Mitchell, S.F. & Parker, R. Principles and properties of eukaryotic mRNPs. Mol Cell 54, 547–58 (2014).

27. Ivanov, P., Kedersha, N. & Anderson, P. Stress Granules and Processing Bodies in Translational Control. Cold Spring Harb Perspect Biol 11(2019).

28. Brengues, M., Teixeira, D. & Parker, R. Movement of eukaryotic mRNAs between polysomes and cytoplasmic processing bodies. Science 310, 486–9 (2005).

29. Kedersha, N. et al. Dynamic shuttling of TIA-1 accompanies the recruitment of mRNA to mammalian stress granules. J Cell Biol 151, 1257–68 (2000).

30. Parker, D.M., Tauber, D. & Parker, R. G3BP1 promotes intermolecular RNA-RNA interactions during RNA condensation. Mol Cell 85, 571–584 e7 (2025).

31. Trussina, I. et al. G3BP-driven RNP granules promote inhibitory RNA-RNA interactions resolved by DDX3X to regulate mRNA translatability. Mol Cell 85, 585–601 e11 (2025).

32. McInerney, G.M. FGDF motif regulation of stress granule formation. DNA Cell Biol 34, 557–60 (2015).

33. Kristensen, O. Crystal structure of the G3BP2 NTF2-like domain in complex with a canonical FGDF motif peptide. Biochem Biophys Res Commun 467, 53–7 (2015).

34. Vognsen, T., Moller, I.R. & Kristensen, O. Crystal structures of the human G3BP1 NTF2-like domain visualize FxFG Nup repeat specificity. PLoS One 8, e80947 (2013).

35. Schulte, T. et al. Combined structural, biochemical and cellular evidence demonstrates that both FGDF motifs in alphavirus nsP3 are required for efficient replication. Open Biol 6(2016).

36. Biswal, M., Lu, J. & Song, J. SARS-CoV-2 Nucleocapsid Protein Targets a Conserved Surface Groove of the NTF2-like Domain of G3BP1. J Mol Biol 434, 167516 (2022).

37. Jumper, J. et al. Highly accurate protein structure prediction with AlphaFold. Nature 596, 583–589 (2021).

38. Varadi, M. et al. AlphaFold Protein Structure Database in 2024: providing structure coverage for over 214 million protein sequences. Nucleic Acids Research 52, D368–D375 (2023).

39. Schulte, T. et al. Caprin-1 binding to the critical stress granule protein G3BP1 is influenced by pH. Open Biol 13, 220369 (2023).

40. Kikhney, A.G., Borges, C.R., Molodenskiy, D.S., Jeffries, C.M. & Svergun, D.I. SASBDB: Towards an automatically curated and validated repository for biological scattering data. Protein Sci 29, 66–75 (2020).

41. Das, S., Lin, Y.H., Vernon, R.M., Forman-Kay, J.D. & Chan, H.S. Comparative roles of charge, pi, and hydrophobic interactions in sequence-dependent phase separation of intrinsically disordered proteins. Proc Natl Acad Sci U S A 117, 28795–28805 (2020).

42. Hazra, M.K. & Levy, Y. Cross-Talk of Cation-pi Interactions with Electrostatic and Aromatic Interactions: A Salt-Dependent Trade-off in Biomolecular Condensates. J Phys Chem Lett 14, 8460–8469 (2023).

43. Bussi, C., et al. Publisher Correction: Stress granules plug and stabilize damaged endolysosomal membranes. Nature 624, E3 (2023).

44. Joyner, R.P. et al. A glucose-starvation response regulates the diffusion of macromolecules. Elife 5(2016).

45. Piper, P.W. Molecular events associated with acquisition of heat tolerance by the yeast Saccharomyces cerevisiae. FEMS Microbiol Rev 11, 339–55 (1993).

46. Felle, H.H. pH: Signal and messenger in plant cells. Plant Biology 3, 577–591 (2001).

47. Franzmann, T.M. et al. Phase separation of a yeast prion protein promotes cellular fitness. Science 359(2018).

48. Peppicelli, S. et al. Acidity and hypoxia of tumor microenvironment, a positive interplay in extracellular vesicle release by tumor cells. Cell Oncol (Dordr) (2024).

49. Bonnet, U., Bingmann, D., Speckmann, E.J. & Wiemann, M. Aging is associated with a mild acidification in neocortical human neurons in vitro. Journal of Neural Transmission 125, 1495–1501 (2018).

50. Forester, B.P. et al. Age-related changes in brain energetics and phospholipid metabolism. NMR Biomed 23, 242–50 (2010).

51. Jin, X. et al. Effects of pH alterations on stress- and aging-induced protein phase separation. Cell Mol Life Sci 79, 380 (2022).

52. Yu, B., Pettitt, B.M. & Iwahara, J. Dynamics of Ionic Interactions at Protein-Nucleic Acid Interfaces. Acc Chem Res 53, 1802–1810 (2020).

53. Dai, Y. et al. Biomolecular condensates regulate cellular electrochemical equilibria. Cell 187, 5951–5966 e18 (2024).

54. Li, M.Z. & Elledge, S.J. SLIC: a method for sequence- and ligation-independent cloning. Methods Mol Biol 852, 51–9 (2012).

55. Cowieson, N.P. et al. Beamline B21: high-throughput small-angle X-ray scattering at Diamond Light Source. J Synchrotron Radiat 27, 1438–1446 (2020).

56. Blanchet, C.E. et al. Versatile sample environments and automation for biological solution X-ray scattering experiments at the P12 beamline (PETRA III, DESY). Journal of Applied Crystallography 48, 431–443 (2015).

57. Panjkovich, A. & Svergun, D.I. CHROMIXS: automatic and interactive analysis of chromatography-coupled small-angle X-ray scattering data. Bioinformatics 34, 1944–1946 (2018).

58. Manalastas-Cantos, K. et al. ATSAS 3.0: expanded functionality and new tools for small-angle scattering data analysis. J Appl Crystallogr 54, 343–355 (2021).

59. Konarev, P.V., Volkov, V.V., Sokolova, A.V., Koch, M.H.J. & Svergun, D.I.:: a Windows PC-based system for small-angle scattering data analysis. Journal of Applied Crystallography 36, 1277–1282 (2003).

60. Svergun, D.I. Determination of the Regularization Parameter in Indirect-Transform Methods Using Perceptual Criteria. Journal of Applied Crystallography 25, 495–503 (1992).

61. Svergun, D.I. Restoring low resolution structure of biological macromolecules from solution scattering using simulated annealing (vol 76, pg 2879, 1999). Biophysical Journal 77, 2896–2896 (1999).

62. Franke, D. & Svergun, D.I. DAMMIF, a program for rapid ab-initio shape determination in small-angle scattering. J Appl Crystallogr 42, 342–346 (2009).

63. Volkov, V.V. & Svergun, D.I. Uniqueness of shape determination in small-angle scattering. Journal of Applied Crystallography 36, 860–864 (2003).

64. Petoukhov, M.V. et al. New developments in the ATSAS program package for small-angle scattering data analysis. J Appl Crystallogr 45, 342–350 (2012).

65. Panjkovich, A. & Svergun, D.I. SASpy: a PyMOL plugin for manipulation and refinement of hybrid models against small angle X-ray scattering data. Bioinformatics 32, 2062–4 (2016).

66. Schindelin, J. et al. Fiji: an open-source platform for biological-image analysis. Nature Methods 9, 676–682 (2012).

