## Supplemental Figures for "Solution architecture of G3BP1 reveals pH-dependent conformational switching underlying liquid-liquid phase separation"

**Appendix Figure S1-S6**

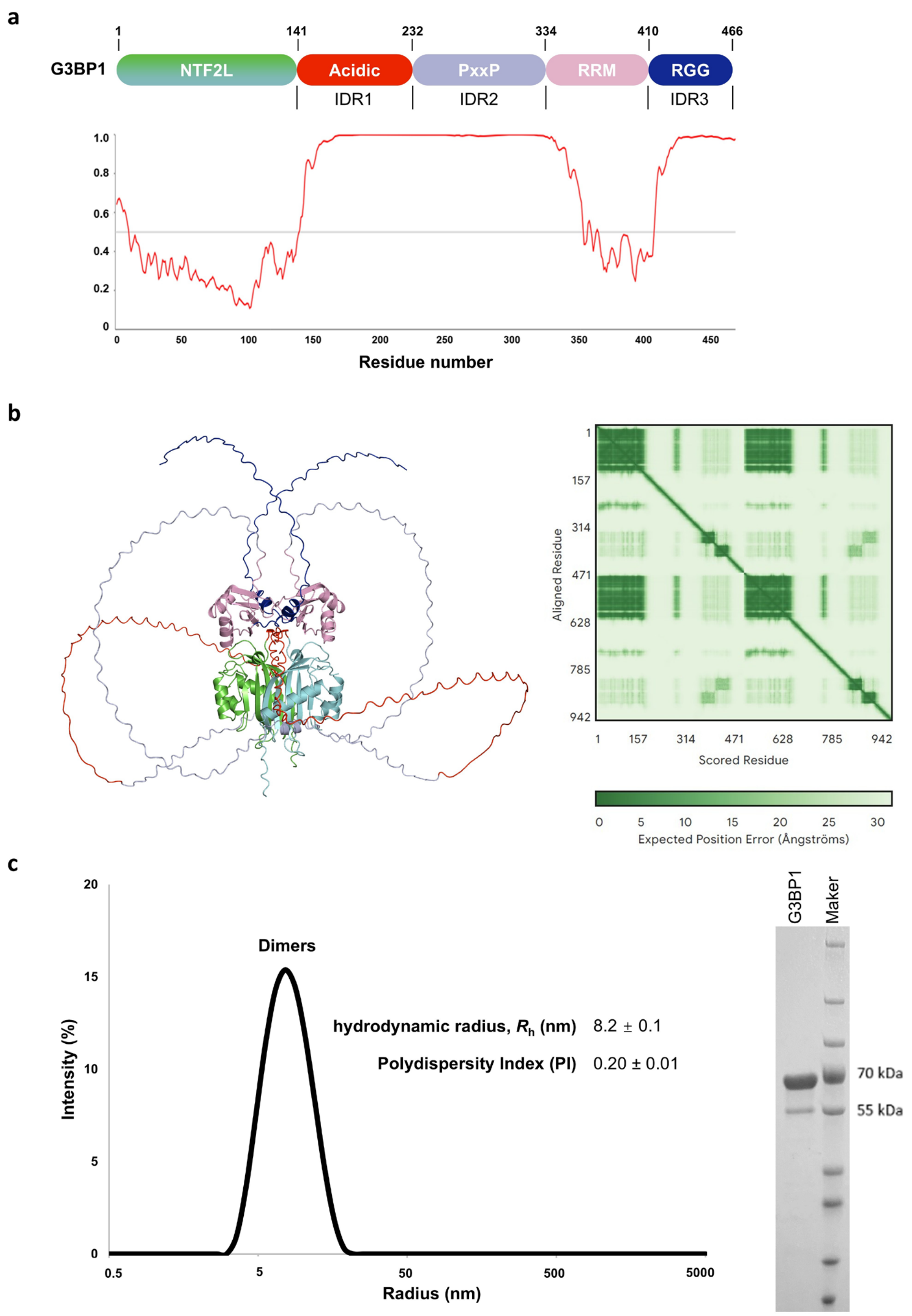


**Appendix Fig. S1: Sequence-based disorder analysis, structural prediction, and purification of full length G3BP1.**

(**a**) Schematic representation of the G3BP1 domain organization aligned with disordered regions predicted using IUPred2A. Scores higher than 0.5 indicate structural disorder. The analysis identifies three major intrinsically disordered regions corresponding to IDR1, IDR2, and IDR3. (**b**) AlphaFold2 structural prediction of the G3BP1 dimer. Left panel: Predicted dimer colored by domain as defined in (a). Right panel: Predicted Aligned Error (PAE) plot, indicating high spatial uncertainty between the NTF2L dimer, RRM, and adjacent IDRs**.** (**c**) DLS measurement of the purified G3BP1 sample exhibits one monodispersed dimer peak at 20 mM Tris pH 7.5 and 150 mM NaCl, with Coomassie blue-stained SDS–PAGE of purified G3BP1. Data represent the mean ± SD of three independent experiments.

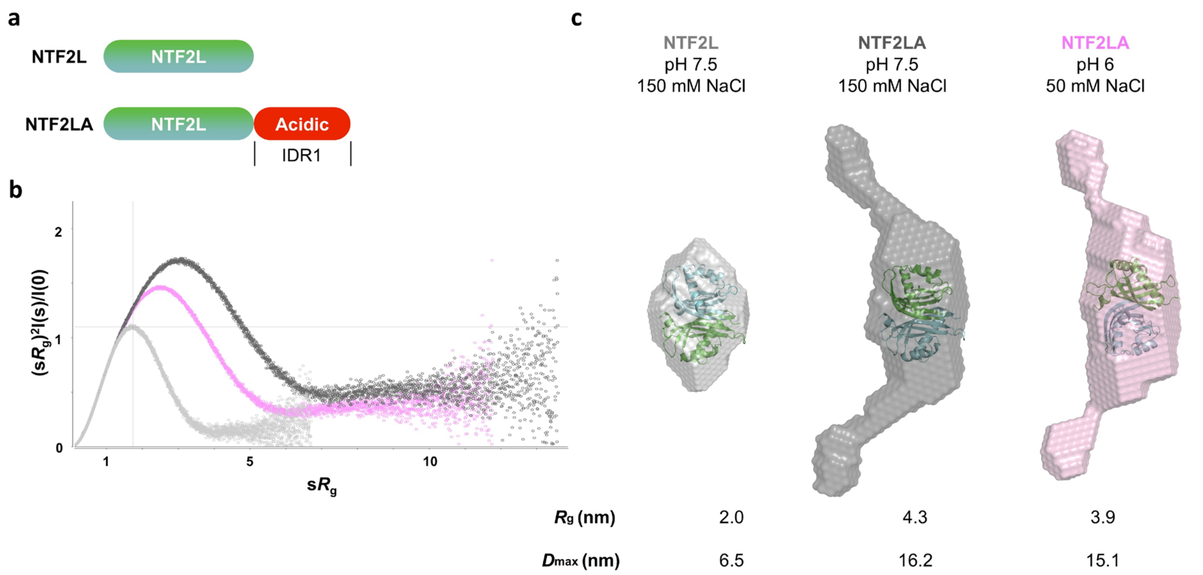


**Appendix Fig. S2: SEC-SAXS analysis of NTF2L and NTF2LA.**

(**a**) Schematic representation of the domain organizations for the NTF2L and the NTF2LA (NTF2L and acidic region) constructs. (**b**) Experimental Scattering data are shown as normalized Kratky plots. NTF2L is shown in light gray (20 mM Tris, 150 mM NaCl, pH 7.5), NTF2LA in dark gray (20 mM Tris, 150 mM NaCl, pH 7.5) and pink (50 mM MES, 50 mM NaCl, pH 6.0). (**c**) The crystal structure of the NTF2L homodimer (PDB code 6TA7) is embedded within each *ab initio* reconstruction (generated with DAMMIF) for NTF2L and the NTF2LA. *Ab initio* models are presented as spheres.

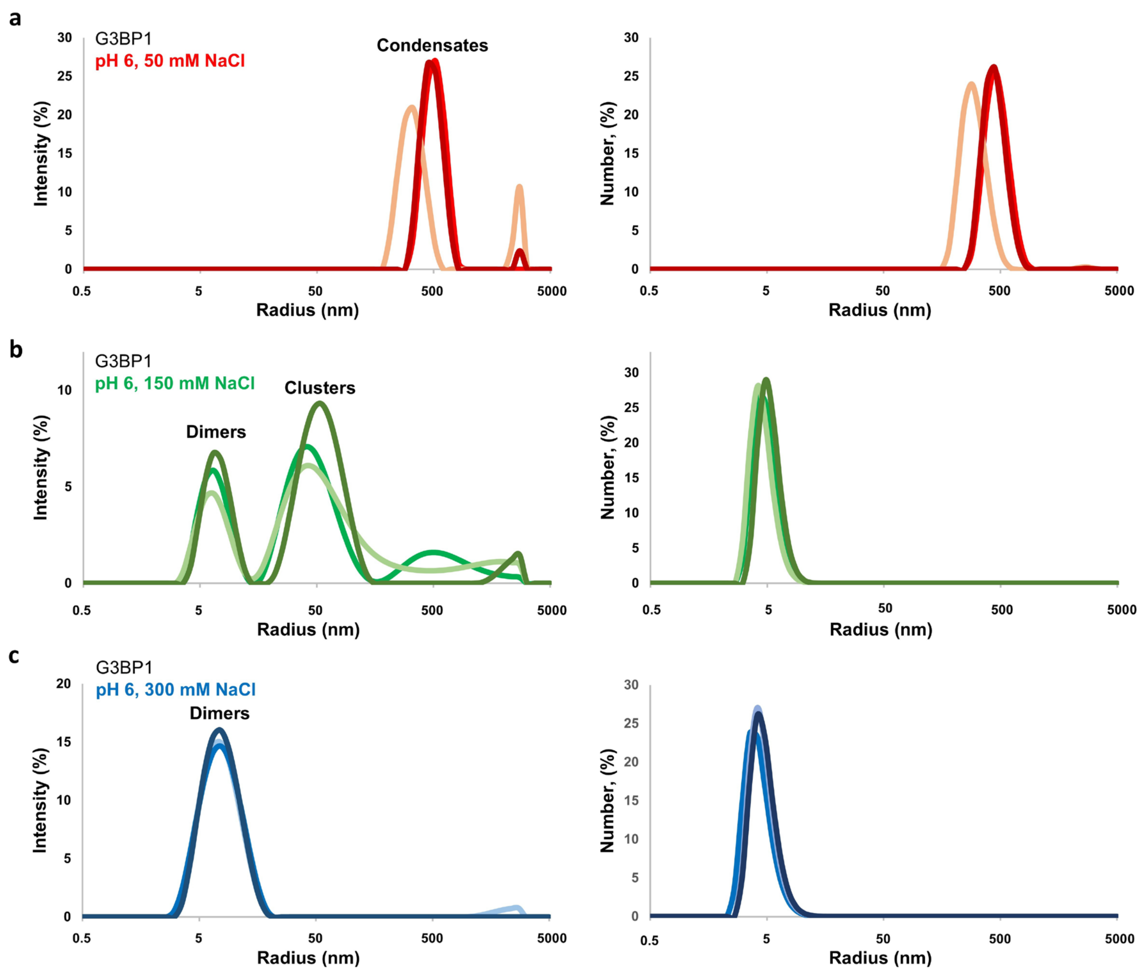


**Appendix Fig. S3: DLS analysis of G3BP1 at pH 6 under varied salt concentrations.**
Particle size distributions of G3BP1 at pH 6.0 in 50 mM (**a**), 150 mM (**b**), and 300 mM (**c**) NaCl. Left and right panels show intensity and number distributions, respectively (n = 3 independent measurements). DLS measurements of the oligomeric species of ΔRGG at pH 6.0 with different salt concentrations. At least two independent DLS measurements were performed.

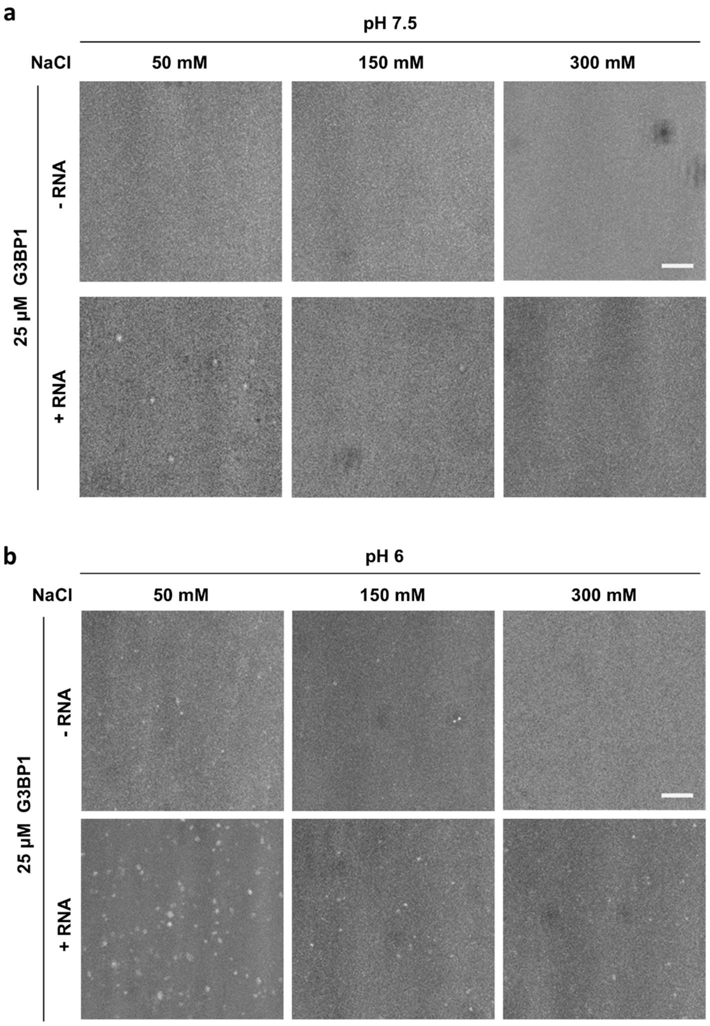


**Appendix Fig. S4: Phase separation of untagged G3BP1 under varying pH and ionic conditions.** (**a**) Phase separation behaviors of untagged G3BP1 (25 µM) with or without 50 ng/µL total RNA at pH 7.5 and the indicated concentrations of NaCl. Scale bar represents 10 µm. (**b**) Phase separation behaviors of untagged G3BP1 (25 µM) with or without total RNA (50 ng/µL) at pH 6 and the indicated concentrations of NaCl. Scale bar represents 10 µm. Phase separation was visualized using a Nikon CrEST X-Light V3 confocal microscope. High-resolution images were acquired using a TL camera. Representative images from at least two independent experiments are shown.

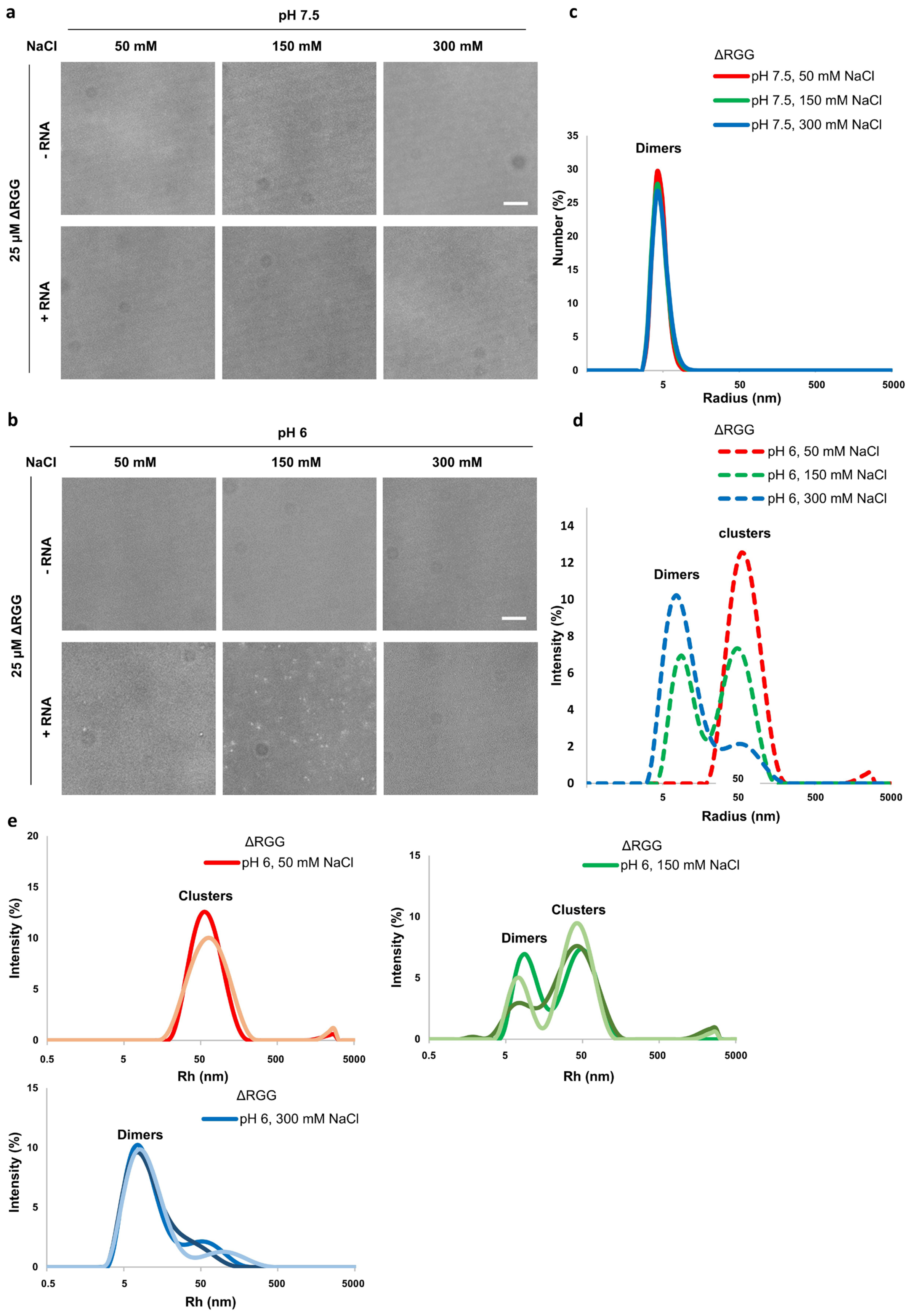


Appendix Fig. S5: Phase separation behaviours and DLS analysis of ΔRGG under varying pH and ionic conditions.

*In vitro* LLPS assay of purified recombinant ΔRGG (25 µM) with or without 50 ng/µL total RNA at pH 7.5 (**a**) and pH 6 (**b**), under indicated salt conditions. Scale bar represents 10 µm. Aligned DLS measurements of the oligomeric species of ΔRGG at pH 7.5 (**c**) and pH 6 (**d**), under indicated salt concentrations, showing the particle size distribution (mean by intensity) profiles. (**e**) DLS measurements of the oligomeric species of ΔRGG at pH 6.0 with different salt concentrations. At least two independent DLS measurements were performed.

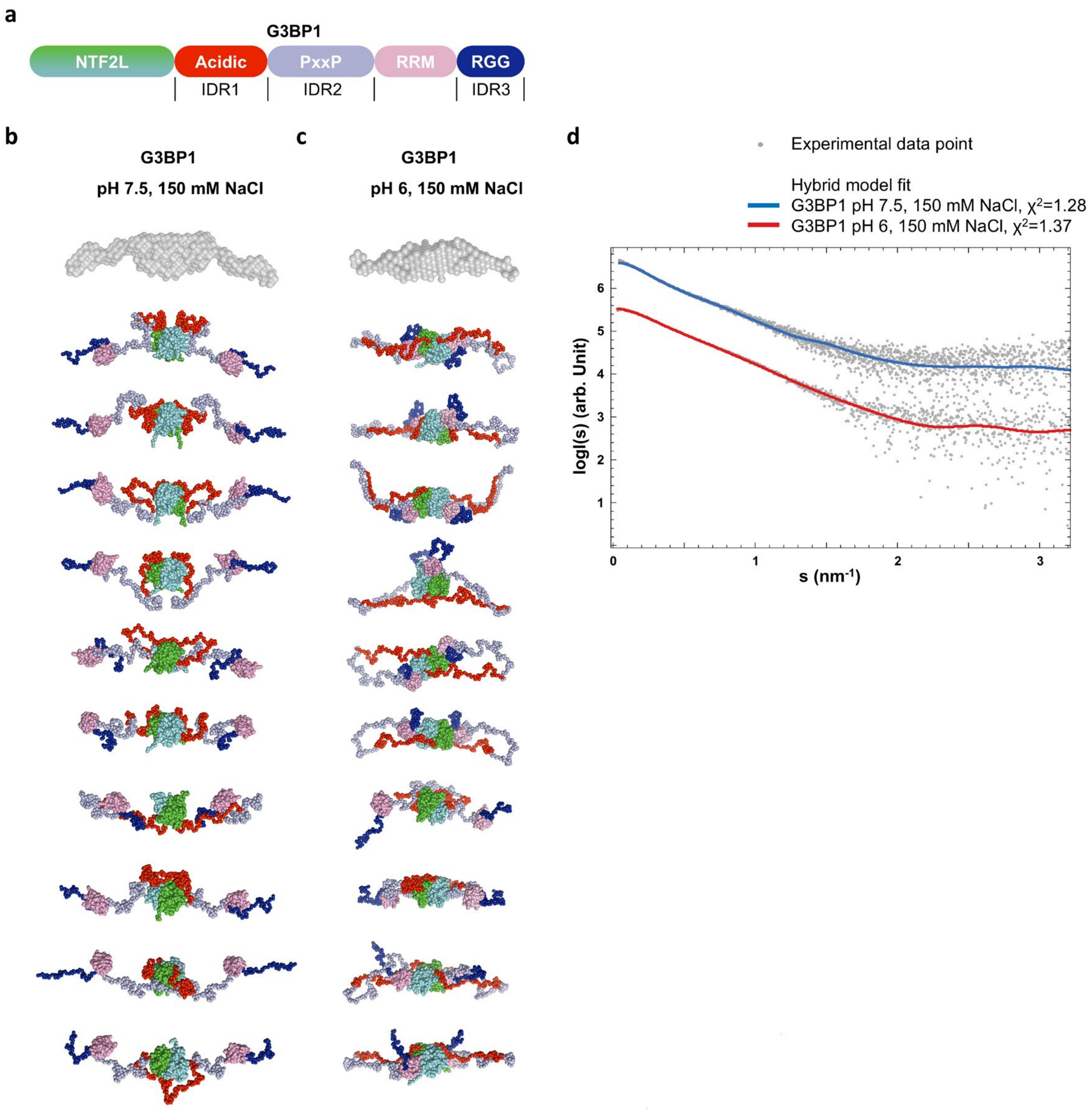


**Appendix Fig. S6: Multiple independent CORAL runs yielded reproducible hybrid models of full-length G3BP1, consistent with the respective overall shapes.** The *ab initio* models (generated with DAMMIN) are represented as transparent surface and the hybrid CORAL models (generated with CORAL) are shown as spheres.

(**a**) Schematic diagram illustrating the domain organization of G3BP1. (**b**) SAXS-based hybrid models of G3BP1 at pH 7.5, 150 mM NaCl are presented, with the domains colored as indicated in (a). The hybrid models feature the centrally localized NTF2L domain and the acidic IDR1. The PxxP, RRM, and RGG domains extend laterally, forming the 'wings' of the structure. (**c**) SAXS-based hybrid models of G3BP1 at pH 6.0, 150 mM NaCl become more compact and the RBD (RRM and RGG region) is positioned closer to the NTF2L homodimer, resulting in a more compact overall shape. (**d**) The fit of representative hybrid models of G3BP1 with experimental scattering data. Curves have been shifted along the y-axis for better visibility.
